# Photosynthetic and water transport strategies of plants along a tropical forest aridity gradient: a test of optimality theory

**DOI:** 10.1101/2023.01.10.523419

**Authors:** Huanyuan Zhang-Zheng, Yadvinder Malhi, Agne Gvozdevaite, Theresa Peprah, Mickey Boakye, Kasia Ziemińska, Stephen Adu-Bredu, Jesús Aguirre-Gutiérrez, David Sandoval, Iain Colin Prentice, Imma Oliveras Menor

**Author notes:** Corresponding author: Huanyuan Zhang-Zheng.

## Abstract

**The research conducted, including the rationale:** The direct effect of aridity on photosynthetic and water-transport strategies is not easy to discern in global analyses because of large-scale correlations between precipitation and temperature. We analyze tree traits collected along an aridity gradient in Ghana, West Africa that shows little temperature variation, in an attempt to disentangle thermal and hydraulic influences on plant traits.

**Methods:** Predictions derived from optimality theory on the variation of key plant traits along the aridity gradient are tested with field measurements.

**results:** Most photosynthetic traits show trends consistent with optimality-theory predictions, including higher photosynthetic capacity in the drier sites, and an association of higher photosynthetic capacity with greater respiration rates and greater water transport. Hydraulic traits show less consistency with theory or global-scale pattern, especially predictions based on xylem efficiency-safety tradeoff. Nonetheless, the link between photosynthesis and water transport still holds: species (predominantly deciduous species found in drier sites) with both higher sapwood-to-leaf area ratio (AS/AL) and potential hydraulic conductivity (Kp), implying higher transpiration, tend to have both higher photosynthetic capacity and lower leaf-internal CO_2_.

**Conclusions:** These results indicate that aridity is an independent driver of spatial patterns of photosynthetic traits, while plants show a diversity of water-transport strategies along the aridity gradient.

*Plain language summary:* Along an aridity gradient in Ghana, West-Africa, we used optimality theory to explain that aridity is an important driver of photosynthetic traits, independent of temperature. Toward drier sites, plants have higher photosynthetic capacities per leaf area but have fewer leaves. We also explain how plants arrange water transportation to support quicker photosynthesis at drier sites. However, plants at the drier sites seem to have diverse combinations of hydraulic traits to satisfy the need for photosynthesis. We reported surprising data-theory inconsistency for some hydraulic traits along the aridity gradient where further research is needed.

## Introduction

Three key photosynthetic processes are frequently considered when seeking to understand plants photosynthesis strategies: light availability and electron transport; aridity and water transport; and CO_2_ concentration and carboxylation (Farquhar et al., 2001). Plants capacities in these photosynthetic processes vary considerably along environmental gradients (Wang *et al*., 2017a; Bahar *et al*., 2017; Yang *et al*., 2019; Oliveras *et al*., 2020). Recently, many efforts have been made to propose universal rules to explain worldwide plant photosynthetic strategies, frequently cited as ‘optimality theories’, which could serve as a basic theoretical framework for vegetation carbon modelling and enable quantitative predictions of key photosynthetic traits (Franklin *et al*., 2020; Harrison *et al*., 2021).

One of the main challenges confronting these universal rules is to explain the ‘pure’ effect of aridity on photosynthesis (Rogers *et al*., 2017). Such challenges become particularly pressing in the context of climate change as greater atmospheric dryness (water vapour deficit, VPD) is predicted for most places (Neelin *et al*., 2006; Grossiord *et al*., 2020a; Bauman *et al*., 2022), which may strongly influence photosynthesis and hence the carbon cycle (Canadell *et al*., 2021). Although optimality theory has shown to successfully explain photosynthetic strategies on multiple scales (Peng *et al*., 2020; Dong *et al*., 2020; Harrison *et al*., 2021), in previous studies, aridity was confounded with temperature, especially when VPD is used as a metric of aridity. Temperature is a stronger driver of photosynthesis than aridity (Smith *et al*., 2019; Peng *et al*., 2021), but few studies have tried to disentangle aridity from temperature (Grossiord *et al*., 2020a). To date, the optimality-theoretical expectation for the impact of aridity on plant traits has not been summarized and tested. Most current earth systems models predict a negative relationship between photosynthesis (denoted by CO_2_ assimilation rate per leaf area, A_area_) andVPD simply due to the closing of stomata without incorporating the dynamics of photosynthetic capacity (denoted by electron-transport capacity, J_max25_ and Rubisco carboxylation capacity standardized to 25 °C, V_cmax25_) (Wang *et al*., 2017a; Green *et al*., 2020). On the contrary, a study focusing on Amazonia argued that photosynthetic capacity is higher for leaves grown in dry season which counteracts the reduced stomatal conductivity, leading to higher photosynthesis under drier climates (Restrepo-Coupe *et al*., 2013; Green *et al*., 2020). Globally higher V_cmax25_ was indeed found for plants grown in drier sites (Cernusak *et al*., 2011; Peng *et al*., 2021; Dong *et al*., 2022). Under experimental conditions, plants grown under low VPD show no difference in CO_2_ assimilation to plants grown under normal VPD (Cunningham, 2005).

Despite the need of incorporating the dynamics of photosynthetic capacity in models, the stand-alone effect of aridity on photosynthesis adaptation still remains unclear. There are two particular challenges. First, aridity can be confounded with temperature on a large spatial scale or temporal scale (Grossiord *et al*., 2020a). Second, optimality theory predicted higher V_cmax_ and A_area_ under higher VPD (Smith *et al*., 2019) but it is unclear how plants in drier environments arrange water transportation through xylem to support higher A_area._ A comprehensive theoretical framework is lacking to incorporate the effect of VPD on all leaf-level photosynthesis processes (light, water and CO_2_) with consideration of water delivery to leaves (Mencuccini *et al*., 2019a).

Here, we examine a dataset of detailed traits measurements along an aridity gradient in West African forests to disentangle the effect of aridity on photosynthesis from temperature and to explain the effect with optimality theory. The key questions we address are: (1) do plants in drier environments have higher photosynthesis rates and how do aridity and photosynthesis interact? (2) If photosynthetic rates are higher in arid environments, as predicted by optimality theory, how do plants arrange greater water transportation under greater atmospheric dryness? To answer these questions, we adopted a theory-data comparison approach where we first review the expectation of recent ‘universal’ theories and deduced 16 testable predictions (some of which have previously been tested but with confounding results). We then examined the consistency between each prediction and field measurement along the aridity gradient (Table 1). Consistency would give field-observed patterns a mechanistic explanation and reinforce the stand-alone impact of aridity on the corresponding trait, while inconsistency would imply weakness of the theory and the possible effect of other environmental factors (like soil properties). Before closing the paper, we summarize the consistency and inconsistency with an integrated theoretical framework to address the ‘pure’ effect of aridity on photosynthesis.

**Table 1.**
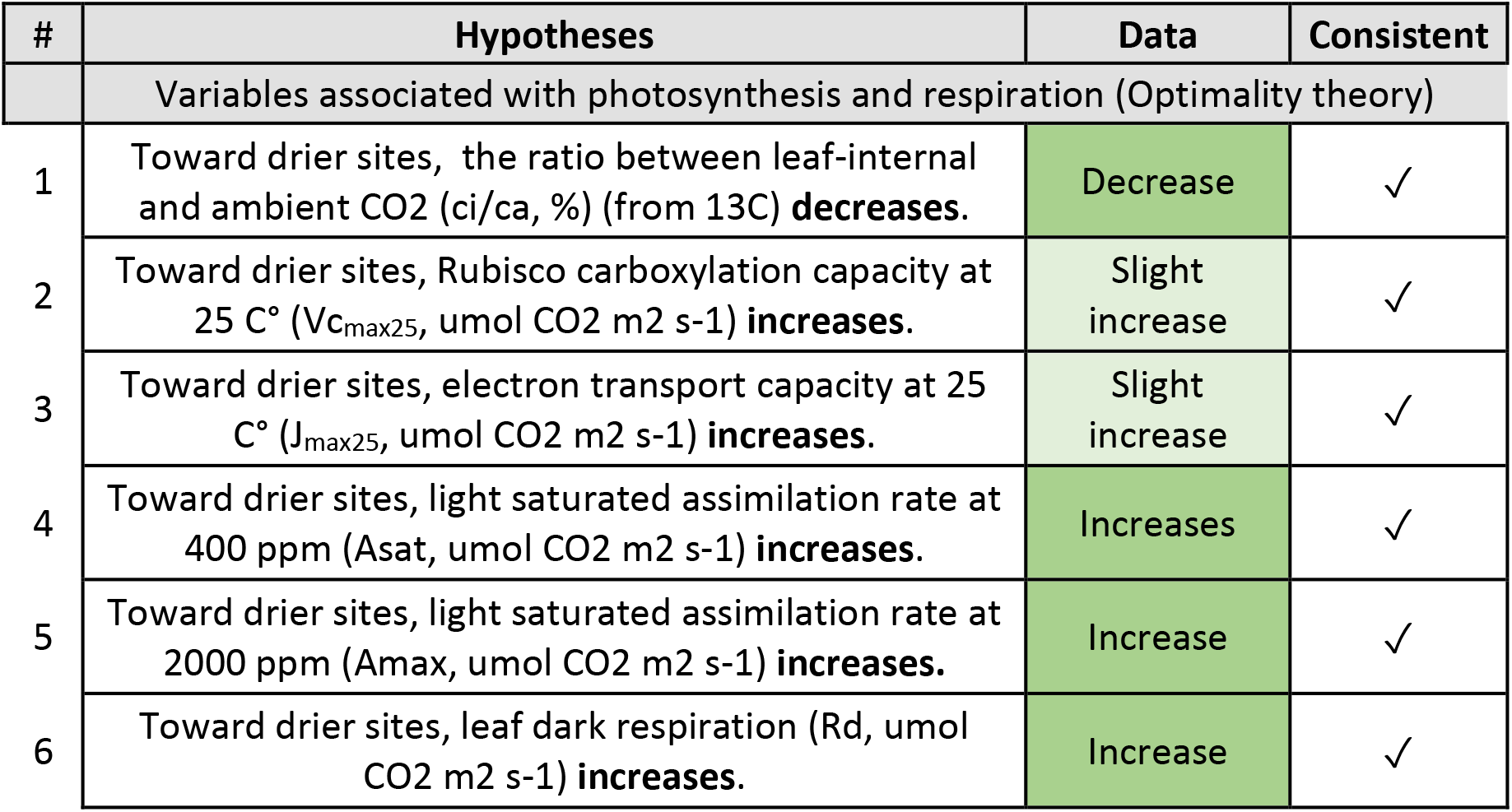
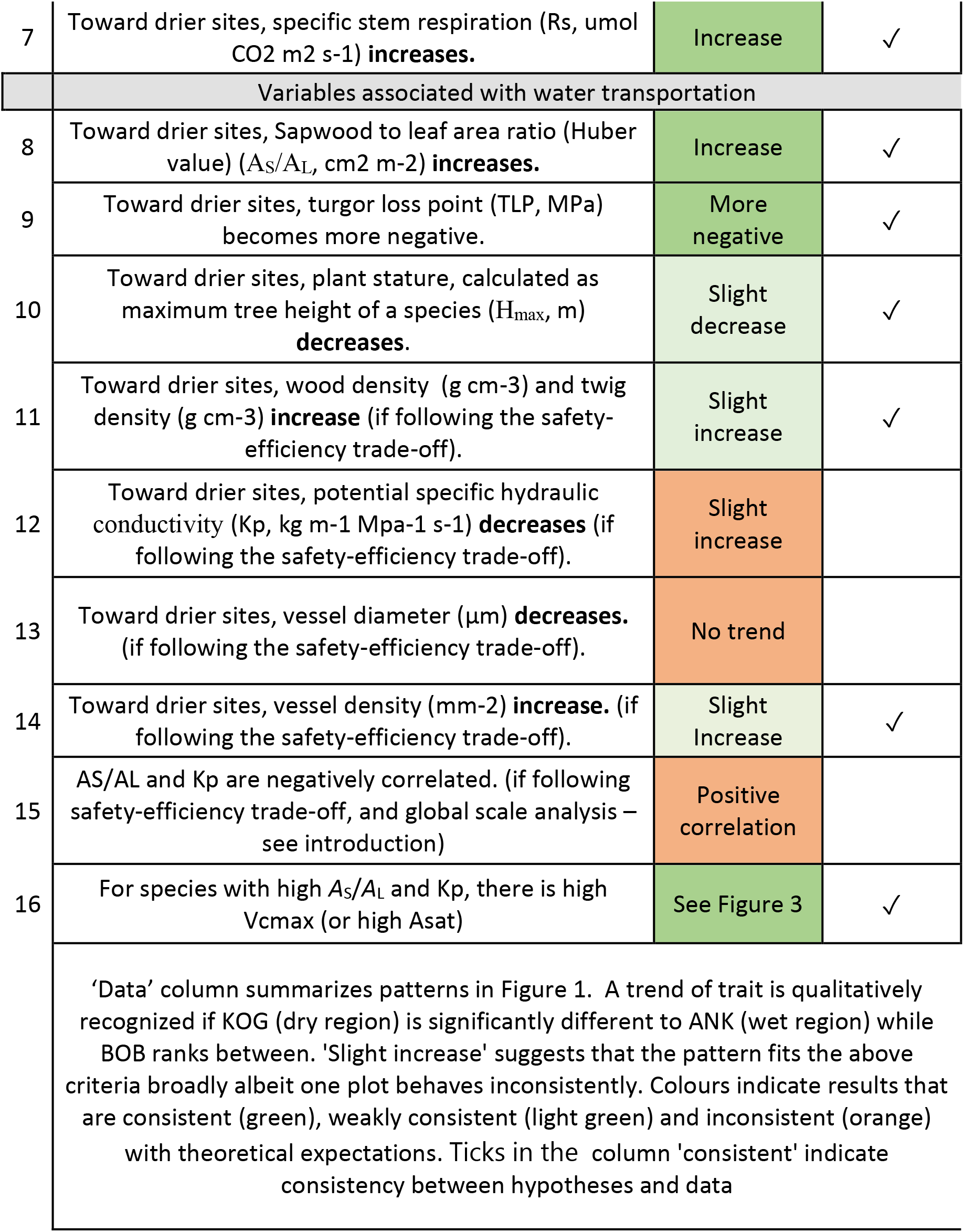
Traits name, unit, hypotheses and findings from field measurements along the rainfall gradient, Green color denotes consistency between theory and our field data. Orange color denotes inconsistency.

| # | Hypotheses | Data | Consistent |
| --- | --- | --- | --- |
| Variables associated with photosynthesis and respiration (Optimality theory) |  |  |  |
| 1 | Toward drier sites, the ratio between leaf-internal and ambient CO<sub>2</sub> (ci/ca, %) (from 13C) **decreases**. | Decrease | ✓ |
| 2 | Toward drier sites, Rubisco carboxylation capacity at 25 C° (V<sub>cmax25</sub>, umol CO<sub>2</sub> m<sup>2</sup> s<sup>-1</sup>) **increases**. | Slight increase | ✓ |
| 3 | Toward drier sites, electron transport capacity at 25 C° (J<sub>max25</sub>, umol CO<sub>2</sub> m<sup>2</sup> s<sup>-1</sup>) **increases**. | Slight increase | ✓ |
| 4 | Toward drier sites, light saturated assimilation rate at 400 ppm (Asat, umol CO<sub>2</sub> m<sup>2</sup> s<sup>-1</sup>) **increases**. | Increases | ✓ |
| 5 | Toward drier sites, light saturated assimilation rate at 2000 ppm (Amax, umol CO<sub>2</sub> m<sup>2</sup> s<sup>-1</sup>) **increases**. | Increase | ✓ |
| 6 | Toward drier sites, leaf dark respiration (Rd, umol CO<sub>2</sub> m<sup>2</sup> s<sup>-1</sup>) **increases**. | Increase | ✓ |
| 7 | Toward drier sites, specific stem respiration ($R_s$, $\mu\text{mol CO}_2 \text{ m}^{-2} \text{ s}^{-1}$) **increases**. | Increase | ✓ |
| Variables associated with water transportation |  |  |  |
| 8 | Toward drier sites, Sapwood to leaf area ratio (Huber value) ($A_s/A_L$, $\text{cm}^2 \text{ m}^{-2}$) **increases**. | Increase | ✓ |
| 9 | Toward drier sites, turgor loss point (TLP, MPa) becomes more negative. | More negative | ✓ |
| 10 | Toward drier sites, plant stature, calculated as maximum tree height of a species ($H_{\text{max}}$, m) **decreases**. | Slight decrease | ✓ |
| 11 | Toward drier sites, wood density ($\text{g cm}^{-3}$) and twig density ($\text{g cm}^{-3}$) **increase** (if following the safety-efficiency trade-off). | Slight increase | ✓ |
| 12 | Toward drier sites, potential specific hydraulic conductivity ($K_p$, $\text{kg m}^{-1} \text{ Mpa}^{-1} \text{ s}^{-1}$) **decreases** (if following the safety-efficiency trade-off). | Slight increase | |
| 13 | Toward drier sites, vessel diameter ($\mu\text{m}$) **decreases**. (if following the safety-efficiency trade-off). | No trend | |
| 14 | Toward drier sites, vessel density ($\text{mm}^{-2}$) **increase**. (if following the safety-efficiency trade-off). | Slight Increase | ✓ |
| 15 | $A_s/A_L$ and $K_p$ are negatively correlated. (if following safety-efficiency trade-off, and global scale analysis – see introduction) | Positive correlation | |
| 16 | For species with high $A_s/A_L$ and $K_p$, there is high $V_{\text{cmax}}$ (or high $A_{\text{sat}}$) | See Figure 3 | ✓ |
| 'Data' column summarizes patterns in Figure 1. A trend of trait is qualitatively recognized if KOG (dry region) is significantly different to ANK (wet region) while BOB ranks between. 'Slight increase' suggests that the pattern fits the above criteria broadly albeit one plot behaves inconsistently. Colours indicate results that are consistent (green), weakly consistent (light green) and inconsistent (orange) with theoretical expectations. Ticks in the column 'consistent' indicate consistency between hypotheses and data |  |  |  |

### Review of Optimality theory

‘Optimality theory’ was developed recently with the assumption that plants can optimize photosynthesis and minimize maintenance costs according to their living environments, which provides a universal explanation of the variation of photosynthetic strategies under different growing environments (Prentice *et al*., 2014; Sperry *et al*., 2017; Wang *et al*., 2017b; Mencuccini *et al*., 2019a; Stocker *et al*., 2020; Xu *et al*., 2021). Although the above-cited studies have tested the theories on global scales and along elevation gradients, discussion and validation of these theories along aridity gradients, are still lacking. Therefore, we first review the implication of such theories on plants photosynthetic strategies along aridity gradients.

As predicted by the ‘least-cost hypothesis’ (Wright *et al*., 2001, 2003; Medlyn *et al*., 2011; Prentice *et al*., 2014), plants in dry climates maximize the carbon return per molecule of water by keeping stomata relatively closed. Thus, in drier sites, plants are expected to have a lower leaf internal-to-external CO_2_ ratio (c_i_/c_a_) and lower stomatal conductance (g_s_). The ‘coordination hypothesis’ (Beerling & Quick, 1995; Maire *et al*., 2012; Walker *et al*., 2014) assumes equilibrium between Rubisco-limited photosynthesis rates (A_C_) (depending on V_cmax25_ and c_i_) and electron transport-limited photosynthesis rates (A_J_) (depending on J_max25_ and leaf absorbed photosynthetic photon flux density, PPFD) (see the quantitative expression in (Wang *et al*., 2017b; Smith *et al*., 2019; Stocker *et al*., 2020)). To maintain such an equilibrium, plants in drier sites are expected to have larger V_cmax25_ to compensate for the lower c_i_. Otherwise, A_C_ would be lower than A_J_ resulting in the waste of light (PPFD). To sum up, lower c_i_ but higher V_cmax25_ is expected toward drier sites if J_max25_ stays constant (in which case A_J_ would be slightly lower due to smaller c_i_).

In reality, toward drier sites, it is common to see higher leaf-absorbed photosynthetic photon flux density (I_abs_) because of less cloud cover and more open canopies. Considering an additional optimality criterion that J_max25_ is acclimated to I_abs_ (Smith *et al*., 2019), supported by multiple experiments (Björkman, 1981; Ögren, 1993), we would expect higher J_max25_ and A_J_ in drier sites, which further encourages higher V_cmax25_ (see above paragraph). Higher J_max25_ would give rise to higher A_J_, implying higher A_C_ following the ‘coordination hypothesis’. All the above would lead to high leaf photosynthetic protein cost in dry sites, hence high leaf dark respiration (Rd), and high transpiration stream maintenance cost (see below for transpiration), hence higher stem respiration per leaf area (R_stem_leaf_) (Prentice *et al*., 2014). Note that R_stem_leaf_ is stem respiration per leaf area, different from the commonly reported stem respiration per stem area (R_stem_stem_). Some of the above predictions have been seen on global scale; for example, higher R_d_ has been found in drier sites (Wright *et al*., 2001; Atkin *et al*., 2015) and higher assimilation rate has been reported from drier sites (Cernusak *et al*., 2011; Maire *et al*., 2015; Peng *et al*., 2021; Dong *et al*., 2022).

It is worth noting that V_cmax25_, g_s_ and c_i_ in this paper are discussed as an overall value for a forest stand, disregarding diurnal variation and intraspecific variation (Stangl *et al*., 2019; Han *et al*., 2022). For instantaneous measurements, there is a positive correlation between A_sat_ (light-saturated assimilation rate at 400 ppm), V_cmax25_, g_s_ and c_i_ (Wright et al., 2003; Fig.2 in Prentice et al., 2014), instead of the opposite trend of V_cmax25_ and c_i_/c_a_ discussed above regarding spatial variation only.

Photosynthesis strategies predicted by the optimality theory above can be linked with stem xylem water transportation strategies via stomatal behaviour, as given by Fick’s law,

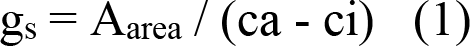

Where g_s_ is stomatal conductance (umol CO_2_ m^-2^ s^-1^), A_area_ is CO_2_ assimilation rate per leaf area (umol CO_2_ m^-2^ s^-1^), and leaf internal (ci, ppm) and external (ca, ppm) CO_2_ concentration

We focus on daytime conditions that produce maximum rates of transpiration and photosynthesis, when water loss through stomata must equal water transport through xylem (assuming no change of stored water) (Brodribb *et al*., 2002; Xu *et al*., 2021):

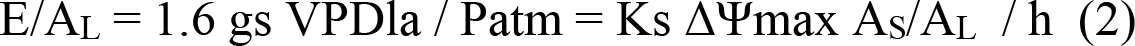

Where E/A_L_ is water transpired per leaf area surface (mol m^-2^ s^-1^), VPDla is leaf-to-air VPD, Patm is atmospheric pressure (Mpa), Ks is sapwood-specific hydraulic conductivity (mol m^−1^s^−1^ MPa^−1^); A_S_/A_L_ is the ratio of sapwood to leaf area (m^2^m^−2^), ΔΨmax is the maximum decrease in water potential from soil to leaves (MPa), h is the transpiration stream path length (m), roughly equivalent to plant height, 1.6 * g_s_ * VPD_la_ / P_atm_ denotes ‘water loss through stomata’, and Ks ΔΨ_max_ A_S_/A_L_ / h denotes water transport through xylem.

Combining the above two equations we obtain a link between water transportation and photosynthesis:

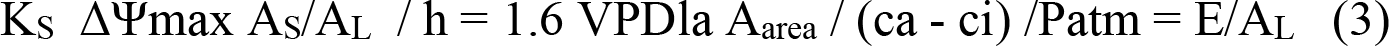

Which could be rearranged to focus on carbon gain:

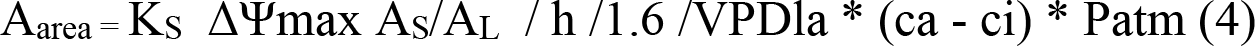

Note that Equation 3 was presented on whole-tree level but was tested using shoot level traits (Xu *et al*., 2021), as well as in this study. Here we disregard diurnal or seasonal variation. Relationships could be very different at other time scales (Mencuccini *et al*., 2019a).

In drier sites with higher VPD, despite smaller g_s_, there should inevitably be a larger E/A_L_ (Granier *et al*., 1996) and more negative ΔΨ_max_ (Gleason *et al*., 2013); therefore smaller maximum tree height (Equation 3), and more negative turgor loss point (TLP, Mpa) in drier sites to increase hydraulic resistance (note that TLP must be more negative than ΔΨmax) (Ryan & Yoder, 1997; Bartlett *et al*., 2012). Equation 3 implies that in drier sites with high VPD, plants require a larger A_S_/A_L_ and/or larger Ks in order to support the same amount of photosynthesis with enhanced transpiration. Following the xylem safety–efficiency trade-off (Manzoni *et al*., 2013; Gleason *et al*., 2016; Bittencourt *et al*., 2016; Grossiord *et al*., 2020b), plants at drier sites would be expected to have lower hydraulic conductivity (K_S_). Although arguments against this trade-off exist (Gleason *et al*., 2016; Körner, 2019; Liu *et al*., 2021), here we present testable hypotheses expected by the trade-off. At dry sites, lower hydraulic conductivity is often associated with smaller vessel diameter, higher vessel density and higher wood density (Poorter *et al*., 2010; Schuldt *et al*., 2013; Hoeber *et al*., 2014). Such patterns have been observed along an Australian aridity gradient (Gleason *et al*., 2013; Pfautsch *et al*., 2016), but no effect of aridity on vessel diameter was reported elsewhere (Olson & Rosell, 2013; Olson *et al*., 2014). Plants in drier sites should have increased hydraulic safety - more negative TLP and more negative P50 (Hacke *et al*., 2001; Martínez-Vilalta *et al*., 2009; Gleason *et al*., 2013; Togashi *et al*., 2015; Liu *et al*., 2019; López *et al*., 2021). In short, toward drier sites, we would expect to see, higher A_S_/A_L_ and more negative TLP. The safety-efficiency trade-off implies lower K_S_, smaller vessel diameter, higher vessel density and higher wood density.

The trade-off between Ks and A_S_/A_L_ is also embedded in the variance of traits in equation 3. K_S_ and A_S_/A_L_ could vary by two orders of magnitude (100-fold variation) (Mencuccini *et al*., 2019b) on a global scale, while ci/ca and A_area_ vary much less (ci/ca: 2 fold; A_area_: 10 fold) (Wright *et al*., 2004; Wang *et al*., 2017b). This leads to a trade-off between K_S_ and A_S_/A_L_ (i.e. K_S_ x A_S_/A_L_ should vary less than either of them). However, given that there are also variations of ci/ca, A_area_, h and ΔΨmax, it is possible that different species range along a spectrum from high A_area_ and E/A_L_ to low A_area_ and E/A_L_ while always satisfying equation 3 (Prentice *et al*., 2014).

In short, the above review leads to hypotheses that plants in drier (normally also brighter) sites tend to develop a photosynthesis strategy with less stomatal conductance and lower ci, stronger photosynthetic capacities (larger V_cmax25_, J_max25_ and A_area_) with more maintenance cost (higher Rd and Rs) and larger transpiration per leaf area which the water transport system would adjust to with higher A_S_/A_L_, lower Ks, lower tree height and more negative TLP. We break the above prediction down into 16 testable hypotheses (Table 1) and test each of them along a forest aridity gradient.

## Materials and Methods

### Study sites - the aridity gradient

This study presents and analyses physiological traits data collected from seven one-hectare forest and savanna plots distributed along a wet-dry gradient across three sites, Ankasa (ANK, moist rainforest), Bobiri (BOB, semi-deciduous forest) and Kogyae (KOG, dry forest and mesic savanna), in Ghana, West Africa (Figure S1, S2) (Moore *et al*., 2018; Oliveras *et al*., 2020), as part of the Global Ecosystem Monitoring (GEM) network (Malhi *et al*., 2021). These sites share very similar mean annual temperature but span a steep gradient of aridity (Figure 1), which provided a “natural laboratory” to disentangle the hydraulic aspect of plant traits variation from temperature. Light increase toward drier sites (Table S 1). There is no seasonal variation in temperature. Two rain seasons (Figure S9 S10) in all study sites occurred in similar months but the total amount of precipitation increases from dry to wet site. Latitude, longitude, number of species and more information are provided in Table S 1.

**Figure 1.**
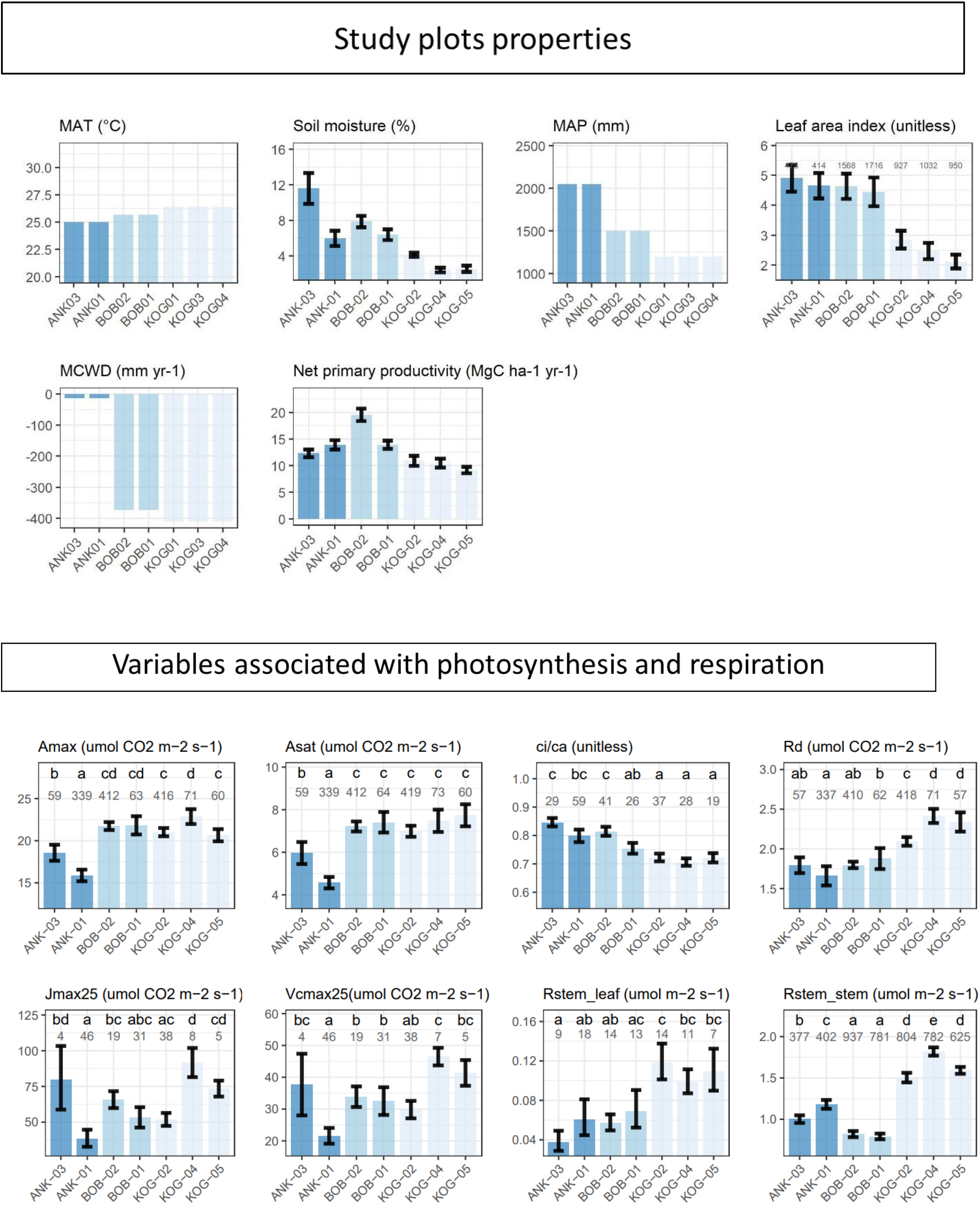
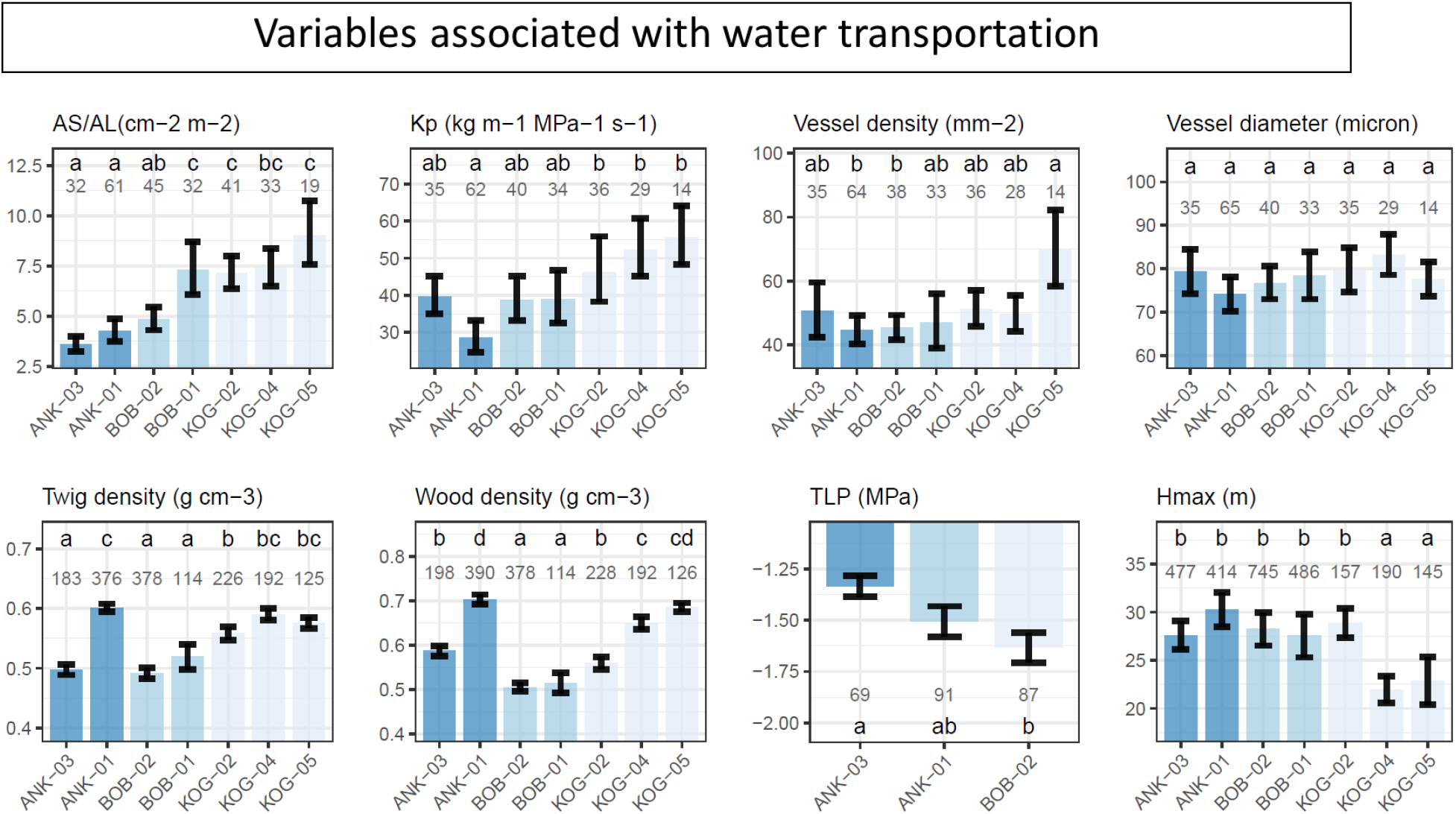
Plot scale community weighted mean (with standard error) from the wettest (left) to the driest (right) plot. Here we show Maximum Cumulative Water Deficit (MCWD), Mean annual air temperature (MAT), mean annual precipitation (MAP) and soil volumetric water content at 12 cm depth (Soil moisture), leaf area index (LAI) and Net primary productivity (NPP) all as annual mean. See Table 1 for the list of plant traits. Forest plots are arrayed from left to right in order of increasing aridity according to MCWD and volumetric water content. The number denotes the number of samples, which could be a leaf, a branch, a tree or a species depending on the variable. The letters denote significance (P<0.05) in plot-to-plot difference.

Although one-hectare plots (e.g. BOB-02) within the same site (e.g. BOB) share very similar air temperature and precipitation, they can differ in terms of belowground water supply due to small-scale variations in soil properties and topography (Table S 1). Along the aridity gradient, there are also variations in soil and vegetation type, with vegetation seasonality and deciduousness increasing considerably towards drier sites. More information about the soil properties and climate of all three sites can be found in (Domingues *et al*., 2010; Chiti *et al*., 2010; Moore *et al*., 2018). Moreover, the swampy rainforest (ANK03) is partly inundated during the wet season but not ANK01, which is located on a hill and never inundated. From KOG02 (dry forest), KOG04 to KOG05 (savanna), forest plots become more deciduous with a smaller number of trees(Table S 1). Nonetheless, many species are ‘semi-decideous’ which shed only part of the canopy in the dry season. Within any site, there are many common species between plots but species composition (e.g., top five abundant species) could still be very different. There are almost no common abundant species between the three sites (ANK, BOB and KOG).

### Aridity indices and soil moisture

Although the Introduction focuses on VPD, we only have one meteorological station at each site which could not tell VPD difference between plots. We thus provide other indices of aridity. At site scales, we provide Maximum Cumulative Water Deficit (MCWD) and Aridity index (the ratio of annual potential evapotranspiration (PET) to mean annual precipitation (MAP)). At plot scale, we reported not only measured surface (12 cm depth) soil volumetric water content, but also hydraulic simulations on plot scales with SPLASH v2.0 (Sandoval & Prentice, 2020). This model requires three sets of input data: (1) field observed climate data at site scale during 2011-2016 (2) soil properties measured following the RAINFOR protocols (Quesada *et al*., 2010); (3) terrain data: root zone was assumed 2m, while upslope drainage area, slope inclination and orientation were extracted from a global dataset (Yamazaki *et al*., 2019). We considered two model output indices: the relative soil moisture saturation (Θ), defined as the volumetric water content (θ) normalized by the volumetric water content at saturation (θ_SAT_); a vegetation water stress index (α), estimated as the ratio of annual actual evapotranspiration (AET) to PET. There are more indices shown in Table S 1 and Table S 2.

### Functional trait data measurements

Leaf traits field campaigns were conducted using a standardized protocol between October 2014 and September 2016 in all plots (Oliveras *et al*., 2020), covering both dry and wet seasons for some traits (see Appendix 1 for sampling protocol). We selected species that contributed to up to 80% of the basal area of each plot and sampled the three largest individuals for each species. From each selected individual, we sampled a sun and a shade branch, and from each branch, we used three leaves and three wood segments to measure leaf and wood traits, respectively. Only sunlit samples were used in this analysis because temperature and light of the shade leaves vary considerably from plot to plot which dilutes the focus on the effect of aridity. The specific number of samples and number of individuals sampled could be found in Table S 1, Figure 1 and Figure S 3.

Wood anatomical traits were analysed in the cross-sectional area of one twig of the sun branch per individual (i.e. three replicates per species) (protocol in Appendix 1). Equation 3 could be interpreted on whole-plant level or shoot level. However, whole-plant traits are challenging to measure and this study is conducted on shoot level. We used K_P_ (potential sapwood-specific hydraulic conductivity) as a proxy of K_S_. K_P_ was calculated from vessel density and vessel diameter following (Poorter *et al*., 2010). Nonetheless, K_P_ and K_S_ may decoupled as not entire sapwwod conducts water (Jacobsen *et al*., 2018). We used twig A_S_/A_L_ as a proxy of whole tree A_S_/A_L_. We used plant stature (H_max_) as a proxy of path length (h). Although H_max_ omits information on root length and multi-layer canopy structure, the proxy would satisfy the need for hypothesis testing in the study region but should be used with caution in future modelling studies.

We calculated stem respiration per leaf area (Rs_leaf) instead of the commonly presented Rs_stem, as a ‘maintenance cost of photosynthesis’ (See Appendix 1) (Prentice *et al*., 2014). To our knowledge, Rs_leaf has not previously been presented with in-situ data in the literature. Here we argue the importance to understand stem respiration from per leaf area perspectives because (1) looking at plans from an integrated view, a leaf does not exists alone but exists associated with a full hydraulic system and Rs_leaf integrates the maintenance cost of this full continuum (Prentice *et al*., 2014) (2) consistency with other photosynthetic traits which were reported per leaf area.

All trait data reported in this study were field-measured except for wood density, which was obtained from a global species database (Zanne *et al*., 2009). Net primary productivity was retrieved from (Moore *et al*., 2018). Global scale sapwood-to-leaf-area ratio in Figure S6 and S4 are sourced from (Mencuccini *et al*., 2019b). Global scale vessel diameter used in Figure S4 is sourced from (Choat *et al*., 2012). Multiple sources of data were joined using species names.

### Statistical analysis

Hypotheses 1-14 (Table 1) were tested by significant differences between wet and dry plots. Principal component analysis (PCA) and standardized major axis regression are used to understand the relationship between Ks, A_S_/A_L_ and photosynthesis traits (Hypothesis 15-16).

We performed a plot-to-plot comparison in answering Hypotheses 1 to 14 as follows: (1) We visually inspected histograms of each trait and transformations to normal distribution were applied if necessary. (2) Outliers were checked with the R package *outliers::scores*, interquartile range method (IQR) with threshold 1.5 (Komsta, 2011). Extreme values were kept when we were sure that they were devoid of errors (3) Community-weighted means were calculated based on the basal area of each species. Standard error was calculated with the same weights (Madansky & Alexander, 2017). (4) Significance of differences in plot-to-plot community-weighted means were then tested with Tukey’s one-way ANOVA using *lm(), glht*(), and *cld*() from *multcomp* package (Hothorn *et al*., 2008), using basal area as weights. In testing Hypotheses 1-14, a hypothesis was accepted if KOG (dry region) was significantly different to ANK (wet region) while BOB (middle aridity) sat in between (Figure 1). (5) Variance partitioning was done with *vegan::varpart()*, following redundancy analysis ordination (RDA) method with the expression: *varpart* (Trait, ∼ Plot, ∼ Species, data = Trait). Variance partitioning reveals whether the change of traits along the aridity gradient was driven by intraspecific or interspecific variation. Note that plots within one site share common species (e.g. ANK01 to ANK03), but species composition is very different between sites (e.g. ANK01 to KOG02). Variance partitioning is also used to diagnose whether the intra-specific variation or measurement errors are overwhelming. To double-check the impact of intraspecific variation, we recalculated a community-weighted mean by assuming that the same species share the same value of trait (i.e. remove intraspecific variation) and extrapolated traits value to forest plots without trait measurements (Appendix 5)

For hypothesis 16, we applied Principal Component Analysis (PCA) with *FactoMineR::PCA*() (Lê *et al*., 2008). A_sat_, K_P_, A_S_/A_L_ and V_cmax25_ were log10 transformed. We avoided standardization by disabling ‘scale.unit’ in function PCA() so that the variance of a trait was reflected by the length of an arrow in Figure 2. For hypothesis 15, the slopes and significance of correlation were calculated by Standard Major Axis Regression (*function smatr::sma*()), commonly used for summarizing the relationship between two plant traits (Wright *et al*., 2005; Warton *et al*., 2012) as it considers uncertainties of both axes. All analyses were done at the species level (i.e. each point in Figure 2 represents a species) to compare with other studies and join among datasets. Hypothesis 15 was also tested at the global scale because Ks was reported to negatively correlate with A_S_/A_L_ but there is no report on the global correlation between Kp and A_S_/A_L_ (Appendix 4).

**Figure 2.**
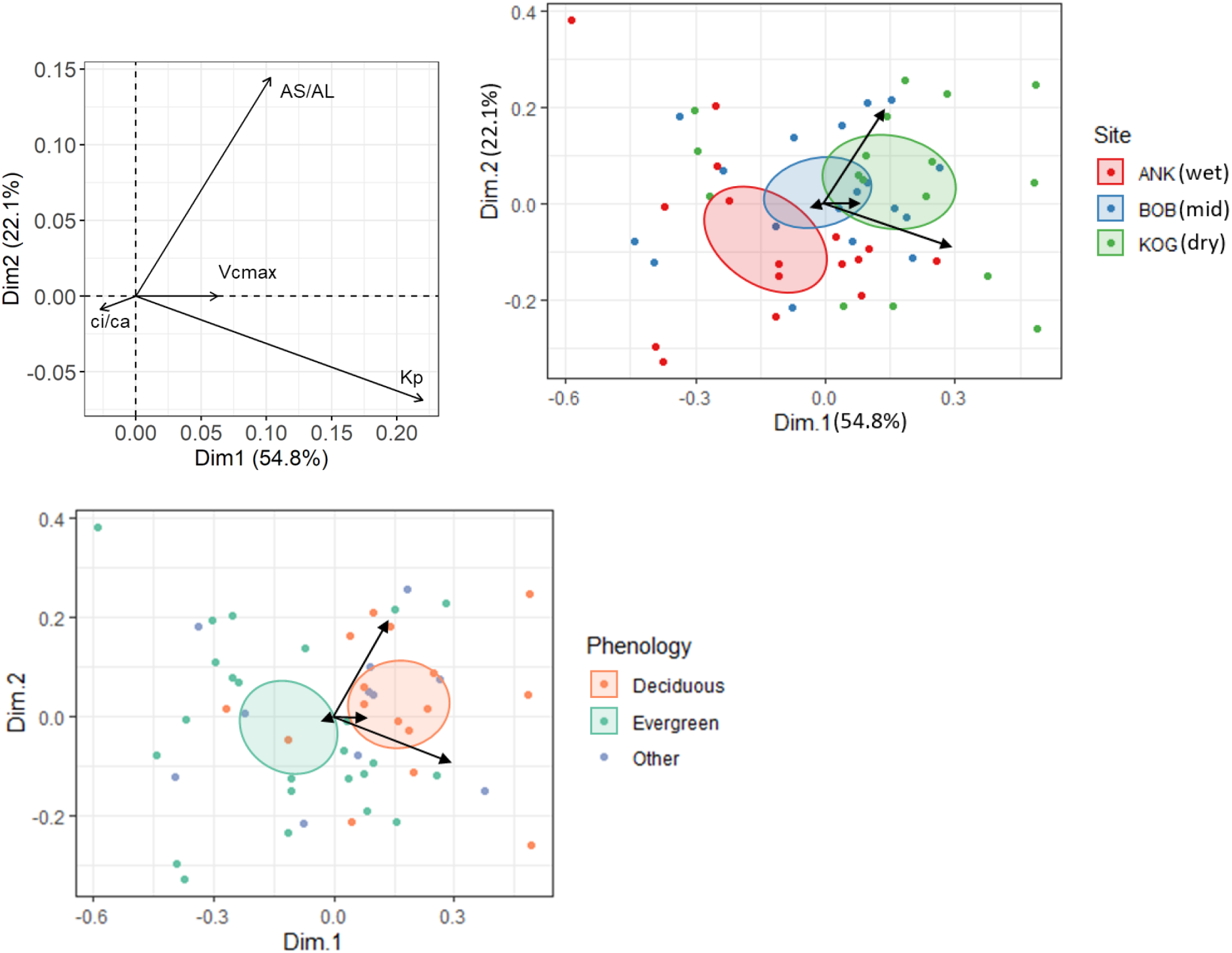
Principal components analysis for Huber value (AS/AL), the ratio between leaf internal and ambient CO2 (ci/ca), Rubisco carboxylation capacity at 25 degree (Vcmax25) and potential specific conductivity (Kp) on species scale. Values are transformed to achieve normal distribution but not standardized to equal variance; therefore the length of arrows denotes the variance of the specific trait. The ellipses for each site are confidence ellipses around group mean points. The PCA axes in all figures are identical. Note that the three figures display the same PCA, but with a different classification of scatter points.

## Results

### Aridity gradient

The values of the aridity index (PET/MAP) (site scale) reveal a clear aridity gradient from ANK (moist rainforest site) to BOB (mid) and KOG (dry) (Table S 2). The same order could be arrived at with VPD or maximum cumulative water deficit (MCWD).

On the other hand, the simulations of relative soil moisture saturation (Θ) and vegetation water stress index (α) (plot scale) show that plants at BOB were the least soil moisture stressed, followed by ANK and KOG. BOB-02 has the highest values in these two metrics, different to surface soil moisture (Figure 1). The model reports the highest runoff at ANK-03, capturing to some degree the seasonal flooding, as also observed in the field. The different patterns of Θ (or α) to aridity index along the aridity gradient are caused by the soil characteristics which in turn define water holding capacity and hydraulic conductivity; for example, the plots in BOB are atmospherically drier (higher PET/MAP) than in ANK but they could hold more water (higher Θ). Especially in BOB-02, the infiltration rate is strongly reduced by low soil saturated hydraulic conductivity (60 mm/hr, less than half of ANK plots), and hence water can stay more time in the root zone while percolating. This acts as a buffer against the evaporative demand, maintaining water availability during dry months. The hydrological modelling outputs also match with field observation of surface soil volumetric water content (Figure 1) and plot vegetation characteristics (Figure 1, S2). For presentation (Figure 1), we rank sites by MCWD and then plots within sites by volumetric water content.

### The effect of aridity on traits

From a photosynthesis perspective, along the aridity gradient, we see consistency between theoretical prediction and field measurements (Table 1) for all traits: toward drier site, ci/ca decreases (0.85 to 0.71), V_cmax25_ increases (21.58 to 46.48 umol CO_2_ m^-2^ s^-1^), J_max25_ increases (38.48 to 91.44 umol CO_2_ m^-2^ s^-1^), R_d_ increases (1.66 to 2.41 umol CO_2_ m^-2^ s^-1^), R_stem_leaf_ increases (0.03 to 0.12 umol CO_2_ m^-2^ s^-1^), A_sat_ increases (4.56 to 7.72 umol CO_2_ m^-2^ s^-1^) and A_max_ increases (15.88 to 22.86 umol CO_2_ m^-2^ s^-1^). The trends of all photosynthetic traits are successfully predicted by theories based on VPD alone (but note that soil moisture and other aridity indices covary with VPD). As leaf economy traits (Figure S3) and soil nutrients (Table S1) overall do not have a clear trend along the gradient, considering nutrient cycling does not seem to aid the prediction of variation of photosynthetic traits along the aridity gradient.

From a water transpiration perspective, the hypotheses are consistent with field measurements for leaf traits. A_S_/A_L_ is higher in drier sites (359.62 to 901.66 cm^2^ m^-2^) and TLP is more negative in drier sites (-1.33 to -1.63 Mpa). However, less consistency is found between theoretical expectations and field measurements for xylem-related traits. Along the aridity gradient, there is an increasing trend of field K_P_ toward drier sites (from 28.62 to 59.29 kg m-1 Mpa^-1^ s^-1^), against the xylem safety-efficiency trade-off. Considering that K_P_ is calculated from vessel diameter and density. Behind the above trend, vessel diameter also contradict the hypotheses. Vessel diameter does not change along the aridity gradient, while vessel density increased toward drier sites (from 44.57 to 69.69 mm^-2^). The theory expects lower K_P_ and hence higher wood density toward drier sites, but the drier plots (KOG04, KOG05) have higher K_P_, higher twig density and higher wood density than the wettest site (ANK03). Meanwhile we also find K_P_ negatively correlates with twig density on species scales (a Simpson’s paradox, see Appendix 4). ANK-01 has very high wood density and twig density which breaks the increasing trend formed by other plots. H_max_ decreases from wet to dry sites as expected.

Using variance partitioning, we find that the plot-to-plot trends of all traits are dominated by inter-specific rather than intra-specific variation (i.e., components [a] are smaller than [b] in Appendix 5) (i.e. the change of species composition). The analogous patterns between twig and wood density along the aridity gradient also support species turnover since twig density was field measured and wood density was parsed from a global database by species (Zanne *et al*., 2009). Nonetheless, variance induced by ‘not changing species and not changing plot’ or simply measurement errors (component [d]) were large for many traits: accounting for 95% of turgor loss point variance, followed by V_cmax25_ (74%) and J_max25_ (66%).

### The coordination between photosynthesis and water transportation

Data from our West African aridity gradient reveal a weak positive correlation between Kp and A_S_/A_L_, contradictory to Hypothesis 15, and inconsistent with the negative correlation that emerged on global scales (Appendix 4). A_S_/A_L_ for the Ghanaian aridity gradient is higher than the pantropical average. For hypothesis 16, we further explore the link between A_S_/A_L_, Kp and photosynthetic trait. Species with both high A_S_/A_L_ and Kp tend to have higher V_cmax25_ and lower ci/ca. Such species tend to be deciduous and appear more in drier plots (Figure 2). There is a larger variance of hydraulic traits compared to photosynthetic traits. The pattern is consistent if we redo the above PCA with Asat instead of V_cmax25_ (Appendix 4). This finding supports hypothesis 16 (Table 1) as well as equation 3.

## Discussion

### The trend of traits along the aridity gradient

Although most hypotheses (Table 1) have been tested with spatially varying aridity at multiple scales (Harrison *et al*., 2021), testing them along the Ghana aridity gradient helps to scrutinize the pattern in the absence of temperature variation. The patterns of all photosynthetic traits measured along the aridity gradient (ci/ca, J_max25_, V_cmax25_, R_d_, R_stem_leaf_, A_sat_, A_max_, namely hypotheses 1-7) are consistent with the theoretical expectations, which underscores that aridity is a direct and critical driver of photosynthetic traits in absence of confounding effect with temperature. The increase of photosynthetic capacity towards drier sites is useful in explaining multiple previous observations: (1) Savanna has higher A_sat_ and A_max_ than wet evergreen forest (Gvozdevaite, 2018; Oliveras *et al*., 2020) (2) In the tropics, drier sites are brighter and warmer where higher photosynthetic capacity imply higher actual CO_2_ assimilation per leaf area. This explains the previous finding that woody savanna has sparse canopy but similar net primary productivity to wet evergreen forest (Figure 1) (Moore *et al*., 2018). As leaf area index decreases toward dry sites and photosynthesis rate increases, the mid-aridity site could be the most productive (Moore *et al*., 2018). (3) For wet Amazonia forests, leaves flushed in dry season have higher photosynthetic capacities which increase forest productivity (Wu *et al*., 2020; Green *et al*., 2020).

From a water transportation perspective, forests in drier sites have higher TLP, lower H_max_ and higher A_S_/A_L_, in support of a greater mid-day transpiration stream (agreed with hypotheses 8-10). However, we found slightly higher K_P_ toward drier sites inconsistent with hypotheses derived from the safety-efficiency trade-off. First, it could be associated with the difference between K_P_ and Ks –vessels embolized in drier sites are not dected by anatomical images and not entire sapwood conducts water (Jacobsen *et al*., 2018). It is possible that the trade-off work well only for single-species studies (Pritzkow *et al*., 2020) and become weak on large scales and across species (Gleason *et al*., 2016; Grossiord *et al*., 2020b). Much higher deciduousness in KOG (dry site) than in the wet sites may play a role as higher hydraulic efficiency was observed from deciduous species or more deciduous forests (Choat *et al*., 2005; Chen *et al*., 2008; Liu *et al*., 2021) as they need less hydraulic safety (Körner, 2019). Furthermore, the increase of K_P_ along the aridity gradient is not inconsistent with previous global analysis: First, the trade-off is not strong (or not a strict 1:1) and per given hydraulic safety, a wide range of efficiency was observed in a global dataset (Gleason *et al*., 2016). Second, for environments with wet soils and dry atmosphere, high hydraulic efficiency was observed which reduce xylem water potentials and thus avoid harmful tension in the first place (Gleason *et al*., 2013). We reported a negative correlation between A_S_/A_L_ and Kp at global scales but a positive correlation along the aridity gradient (Appendix 4). One of the reasons for these contrasting opposite correlations may lie in a geographical sampling bias – the global dataset with scarce data points from West Africa compared with the Ghanaian dataset. The other possibility could be a confounding effect by temperature or vegetation type at the global scale; for example, a negative correlation between A_S_/A_L_ and Ks was reported globally (Mencuccini *et al*., 2019b) and on continental (Australia) scales (Gleason *et al*., 2012), but an insignificant correlation was also reported for tropical forest stands on local scales without varying temperature (Poorter *et al*., 2010; Schuldt *et al*., 2013; Hoeber *et al*., 2014).

By assuming that traits with a clear and strong trend along the aridity gradient are more tightly bound with aridity (Figure 1), ci/ca (stomata behaviour), TLP (drought tolerance) and A_S_/A_L_ (water delivery) are found to be the most aridity-driven traits. The runners-up are R_d_, R_stem_leaf_, J_max25,_ and V_cmax25_, which was thought acclimated to ci/ca and light intensity (Wang *et al*., 2017b). Although ci/ca, V_cmax25_, K_P_ and A_S_/A_L_ all vary from wet to dry sites, we further illustrate that, surprisingly, it is photosynthetic traits instead of hydraulic traits that contrast species from wet to dry sites (also from evergreen to deciduous) (Figure 2). Given that large photosynthetic traits variation from wet to dry plot was induced by species turnover (Appendix 5), our studies hint that facing a drier climate, if allowed time, West African forests photosynthesis could adapt to a drier climate by changing species abundance with possibly more deciduousness and higher photosynthesis capacity albeit less stomatal openness (Aguirre-Gutiérrez *et al*., 2019). Without consideration of the positive effect of aridity on photosynthetic capacity, models could underestimate forest productivity under future drier climates.

### Combining photosynthesis and hydraulic hypotheses

Overall, optimality theory can well explain plant photosynthesis strategies along the aridity gradient and we also expand the theory to consider water transportation. Namely, species in drier sites (with more deciduousness) tend to develop a photosynthesis strategy with less stomata openness (ci/ca), stronger photosynthetic capacities (J_max25_ and V_cmax25_) with more maintenance cost (higher R_d_ and R_stem_leaf_), quicker photosynthesis rate (A_sat_) and larger maximum transpiration per leaf area, supported by larger K_P_ and larger A_S_/A_L_. The product of A_S_/A_L_ and K_P_ is a proxy of water delivery per leaf area, which was previously found well correlated with proxies of photosynthesis rate: Asat (Santiago *et al*., 2004), the quantum yield of electron transport (Brodribb & Feild, 2000) and electron transfer rate (Brodribb *et al*., 2002), consistent with this study (Figure 2). The large variance of wood traits (way larger than leaf photosynthetic traits) (Mencuccini *et al*., 2019b) (Figure 2), hints that plants might have a wide range of choices of traits combinations to provide adequate water transportation (Sperry *et al*., 2002; Prentice *et al*., 2014) in drier sites to support faster photosynthesis. The study also highlights the central role of A_S_/A_L_, or LAI on forest stand scale, or deciduousness on temporal scale, in controlling water relations. Further investigations into xylem functioning are required to further understand how larger water transportation was achieved in drier sites. Although we successfully predicted plants photosynthesis strategies along the aridity gradient (hypothesis 1-7) based solely on VPD without mentioning soil nutrients nor soil moisture, the theoretical deduction implicitly assumes that plants in our study site have adequate access to soil nutrient and belowground water. Further, since soil moisture and VPD co-vary along the aridity gradient and both can cause stomata closure (Rodriguez-Dominguez & Brodribb, 2020), their effects are confounded in this study. Thus caveats should be given to model the effect of aridity using only VPD, especially since soil moisture may be playing a role at other temporal scales (e.g., daily) (Liu *et al*., 2020; Fu *et al*., 2022) or under extreme soil drought (Sperry *et al*., 2002).

### Conclusion

Along the aridity gradient, we find that at drier and brighter sites with more decideousness, species tend to have higher V_cmax25_ and lower ci/ca with both higher A_S_/A_L_ and K_P_ (greater mid-day transpiration steam). With such a working example in West Africa, the study not only underscores the importance of incorporating the positive effect of aridity on photosynthesis capacity, as predicted by optimality theory, in carbon modelling but also explains how plants arrange water transportation for higher photosynthesis at drier sites. The study also highlights the pivotal role of A_S_/A_L_ in plants long-term adjustment to water shortage.

### Data availability

Figures could be downloaded from https://github.com/Hzhang-ouce/Ghana_rainfall_trait_variation_optimality_github. To reproduce figures, data and R codes mentioned in the main text could also be found in the above repository.

## Acknowledgements

We thank Nicolas Raab, Natascha Luijken, Ya-Jun Chen, Yu-Heng Sun, Maurizio Mencuccini, Akwasi DGyamfi, Guillaume Delhaye, Dong Ning, Han Wang, Roberto Salguero-Gómez and Sophie Fauset for valuable discussion and assistance with data processing. Y.M. is supported by the Jackson Foundation. H.Z. received Henfrey Scholarship (from St Catherine’s College, Oxford) and Tang Scholarship (by China-Oxford Scholarship Fund). The field data collection was funded by a grant award (ERC GEM-TRAIT, grant no. ERC-2012-ADG_20120216) to Y.M., with additional support for the fieldwork from a Royal Society-Leverhulme Africa Capacity Building Award and Marie Curie Fellowship to I.O. (FP7-2012-IEF-327990-TipTropTrans).

## Author contributions

IO, YM, ICP, and HZ designed the research and interpreted the results. DS did the hydraulic modelling. YM, AG, TP, MB, KZ, SAB, JAG, IO, HZ contributed to data collection. HZ carried out the analyses and wrote the paper with inputs and revisions from all co-authors.

## Appendix 1 Field sampling protocol

Please note that the number of samples could be found in Table S 1, Figure 1 and Figure S 3 and thus not repeated here.

The ratio between leaf-internal and ambient CO2, ci/ca (unitless) was estimated from leaf δ13C measurements (the stable isotope ratio relative to a standard material). We first estimated Δ13C the difference between the leaf stable isotope ratio and the atmospheric stable isotope ratio, from δ13C at that place and time according to (Cornwell *et al*., 2016). Then we estimated ci/ca from Δ13C by equation 11 in (Peng *et al*., 2020).

Climate and soil variables presented in Figure 1 and Table S1 are field measurements, most of which were sourced from (Moore *et al*., 2018; Oliveras *et al*., 2020). Climate variables were recorded by local meteorological station from 2011 to 2015. Soil properties are average across 0-30cm, field measured in 2013 and 2014. Soil volumetric water content (vwc) (%) was measured in the field every month in 2016, using a soil moisture sensor probe over the depth in the forest over the depth 0–12 cm. Soil hydraulic data in Table S2 are model outputs (See Method).

For light saturated assimilation rate at 400 ppm, Asat (umol CO_2_ m-2 s-1) and at 2000ppm, Amax (umol CO_2_ m-2 s-1), The branch that had been cut was promptly placed in water and recut. To measure leaf gas exchange traits, an open flow gas exchange system (LI-6400XT, Li-Cor Inc., Lincoln, NE, USA) was used. Three leaves were selected from each tree and analyzed for Asat and Amax, as well as dark respiration (R_d_) (µmol m–2 s–1 for all photosynthesis traits). The photosynthetic photon flux density was set at 2000 µmol m-2 s-1, with the exception of dark respiration measurements (0 µmol m-2 s-1). The block temperature was kept constant at 30° C throughout the sampling period, which was similar to the ambient air temperature. More information could be found from supporting information in (Aguirre-Gutiérrez *et al*., 2019; Oliveras *et al*., 2020). The above traits were sampled each month from October 2014 to September 2016.

To determine leaf mass per area, LMA (m-2 kg-1), nitrogen content by area, N_area_ (g m-2), nitrogen content by mass, N_mass_ (g/kg), phosphorus content by area, P_area_ (g m-2), and phosphorus content by mass, P_mass_ (g/kg), we selected three fully-grown trees that emerged from the canopy (total of 298 trees) for each species in a given site. Within each tree, we randomly chose three mature leaves from a fully sunlit branch that were not in the process of senescing. The leaves were then dried in an oven at 70°C until a constant mass was reached. Total leaf lamina area (cm2) was calculated by scanning images using NIH ImageJ (http://rsbweb.nih.gov/ij/) and a custom MATLAB script (https://github.com/bblonder/leafarea). LMA was calculated by dividing the dried leaf mass by the leaf area. Part of these data were reported in (Gvozdevaite, 2018; Oliveras *et al*., 2020). Data reported in this study are slightly different to the above two studies because there is more sampling in this study. Samples were taken each month from October 2014 to September 2016.

To measure maximum rate of electron transport at 25 °C J_max25_ (umol CO2 m-2 s-1) and maximum rate of carboxylation at 25 °C V_cmax25_ (umol CO2 m-2 s-1), we sampled one individual tree per species within each study plot to generate A-Ci curves, which show the photosynthetic response to changes in substomatal CO2 concentration (Ci). CO2 concentration was changed in the following sequence: 400, 300, 200, 100, 50, 400, 600, 800, 1200, 1500, and 2000 µmol m-2 s -1. The photosynthetic photon flux density was set at 2000 µmol m-2 s-1, and the block temperature was kept constant and closest to ambient throughout the sampling period at 30°C. We used the A-Ci curve fitting method and followed the procedure described in detail in Appendix B of Domingues et al. (2010) which extracts V_cmax_ and J_max_ values. To enable comparison of our data and findings with the wider literature on photosynthetic capacity variability, we scaled the measured and estimated values of V_cmax_ and J_max_ to a reference temperature of 25°C following Sharkey et al. (2007). We further refer to the scaled values as V_cmax25_ and J_max25_ in the text. V_cmax25_ and J_max25_ are collected in October 2015. The field campaign did not finish at site ANK-03 leading to very little number of samples at this site.

To calculate sapwood area to leaf area, or Huber value, AS/AL (cm-2 m-2), we first determine leaf area (AL) of a sampled terminal, sun-exposed shoots from the outer canopy. We scanned the adaxial side of the leaf lamina (without petiole) on a Canon Lided220® flatbed scanner and analysed the images using a Matlab code that can be found at https://github.com/bblonder/leafarea Neyret et al. (2016). The total leaf area per branch (AS/AL) was determined with the assumption that the branch diameter (without bark) corresponded to sapwood area. Samples are collected in October 2015.

For structural traits, including twig density (g cm-3), vessel density (mm-2), average vessel diameter (μm) and potential specific hydraulic conductivity, Kp (kg m-1 MPa-1 s-1), we sampled ∼8-10mm diameter twigs, on three replicates per species (Gvozdevaite, 2018). Cross sections about 20-50μm were made using a sliding microtome, stained in safranin O and alcian blue, and permanently mounted on a microscopic slide. A pie shaped segment stretching from pith to cambium was photographed using OptronicsMicrofire camera mounted on Olympus BX-50 microscope and PictureFrame software. All vessels within a pie region were marked and coloured using magic wand tool (GIMP, http://gimp.org) and interactive pen display (Wacom Cintiq 22HD). Vessel area, average diameter (average of minimum and maximum diameters) and pie region area were measured using ImageJ. Next, average diameters of all vessels per given pie area were averaged resulting in sample average vessel diameter (VD) which was then used in the analyses. Vessel density (ρV) was calculated by dividing the total number of vessels in an analyzed pie section by the area of that pie section. Lastly, we calculated Kp using the Hagen-Poiseuille equation as per Poorter et al. (2010). Twigs were dried for at least 72 hours at 105°C and mass was measured on a precision balance. The twig density was calculated as dry mass divided by the volume of soaked wood. Samples are collected in October 2015.

For maximum tree height of a species, Hmax (m), in January 2020, we measured the tree height of each tree using a digital clinometer in the forest plot (with diameter at breast height larger than 10cm).

For turgor loss point, TLP (MPa), a sunlit branch with fully grown leaves was collected and re-cut while submerged in water. The branch was then rehydrated overnight, covered with a black plastic bag sprinkled with water on the inside, and left to rehydrate for approximately 15 hours. To generate pressure-volumn curves, paired measurements of leaf water potential and leaf mass were repeatedly taken after intervals of bench drying, resulting in 9 to 12 points per curve. The leaves were scanned to obtain their area and then oven-dried at 60 degrees Celsius for 3 days to obtain their dry mass. PV curves were fitted to extract TLP using codes in (Raab, 2020). TLP was collected in August 2019. Pressure-volume curves are measured in October 2015.

Wood density was provided by Forestplot.net who sourced information from (Zanne *et al*., 2009). In Appendix 4, we also compared data collected from Ghana with AS/AL from (Mencuccini *et al*., 2019b)and vessel diameter from xylem functional traits database (Choat *et al*., 2012).

Stem respiration per steam area (Rs_stem) was measured using a closed dynamic chamber method, from 25 trees distributed evenly throughout each plot at 1.3 m height with an IRGA (EGM-4) and soil respiration chamber (SRC-1) connected to a permanent collar. As we know tree height of each tree, Rs_stem could be converted to stem respiration per leaf area (Rs_leaf) using tree height and AS/AL. Assuming trees have a cylindrical shape, we have Rs_leaf= Rs_stem *4 * AS/AL * H / DBH, where AS/AL is Huber value, H is tree height and DBH is diameter at breast height. We calculated Rs_leaf because most of the traits and theories involved in this study were expressed on per leaf area basis. Stem respiration was measured every three months from 2014 to 2016.

Leaf area index (LAI) was estimated from hemispherical images taken with a Nikon 5100 camera and Nikon Fisheye Converter FC-E8 0.21x JAPAN near the center of each of the 25 subplots in each plot in each site, at a standard height of 1 m, and during overcast conditions. 22,000 photos were collected in total, every month during 2016-2017(ANK), 2012-2017 (BOB&KOG). Photos were processed using machine learning-based software ‘ilastik’ (Berg *et al*., 2019) for pixel classification and CANEYE (Demarez *et al*., 2008) for leaf area index calculations. The exposure procedure followed (Zhang *et al*., 2005) and GEM manual (Malhi *et al*., 2021) (http://gem.tropicalforests.ox.ac.uk). The following parameters were supplied to CANEYE.

1. P1 = angle of view of the fish eye divided by the amount of pixels from centroid of the fish eye circle to where horizon is on the image.
2. angle of view = 90 degree, in which case, the edge of the photo is the horizon and the centroid of the image is zenith.
3. COI = 80, consideration of field is 80 degrees, we don’t want the edge of the photo because it is not clear and sometimes obscure by tall grasses or saplings.
4. Sub sample factor =1
5. Fcover = 20 degree, this is to calculate the percentage of black pixels within central 20-degree ring. We used this to understand the relative openness of canopy for the given image. It is not relevant to LAI
6. PAIsat = 10, When a pixel is completely black, mathematically, the leaf area index (LAI) is infinite. As we provide CANEYE 25 subplot images for each estimation of LAI, this means all 25 subplot images show black at a given pixel. To address this ‘infinite’issue, we use a value of 10 for LAI in such cases. This value is based on the guess that, the densest point in a tropical forest should have an LAI of 10.
7. Latitude 0 and Day of year a random number (not relevant for tropical site LAI)

Then, we extract output from CANEYE using software R. We chose the latest method of LAI calculation offered by CANEYE, ‘CE V6.1 True PAI’. CANEYE reported one LAI value per method (4 methods) per plot per site per month, as a synthesis across 25 subplots images. As systematic error is dominating in LAI calculation, we take the standard deviation of LAI across four methods as the uncertainty for LAI.

## Appendix 2 Study Sites, the aridity gradient

**Figure S 1.**
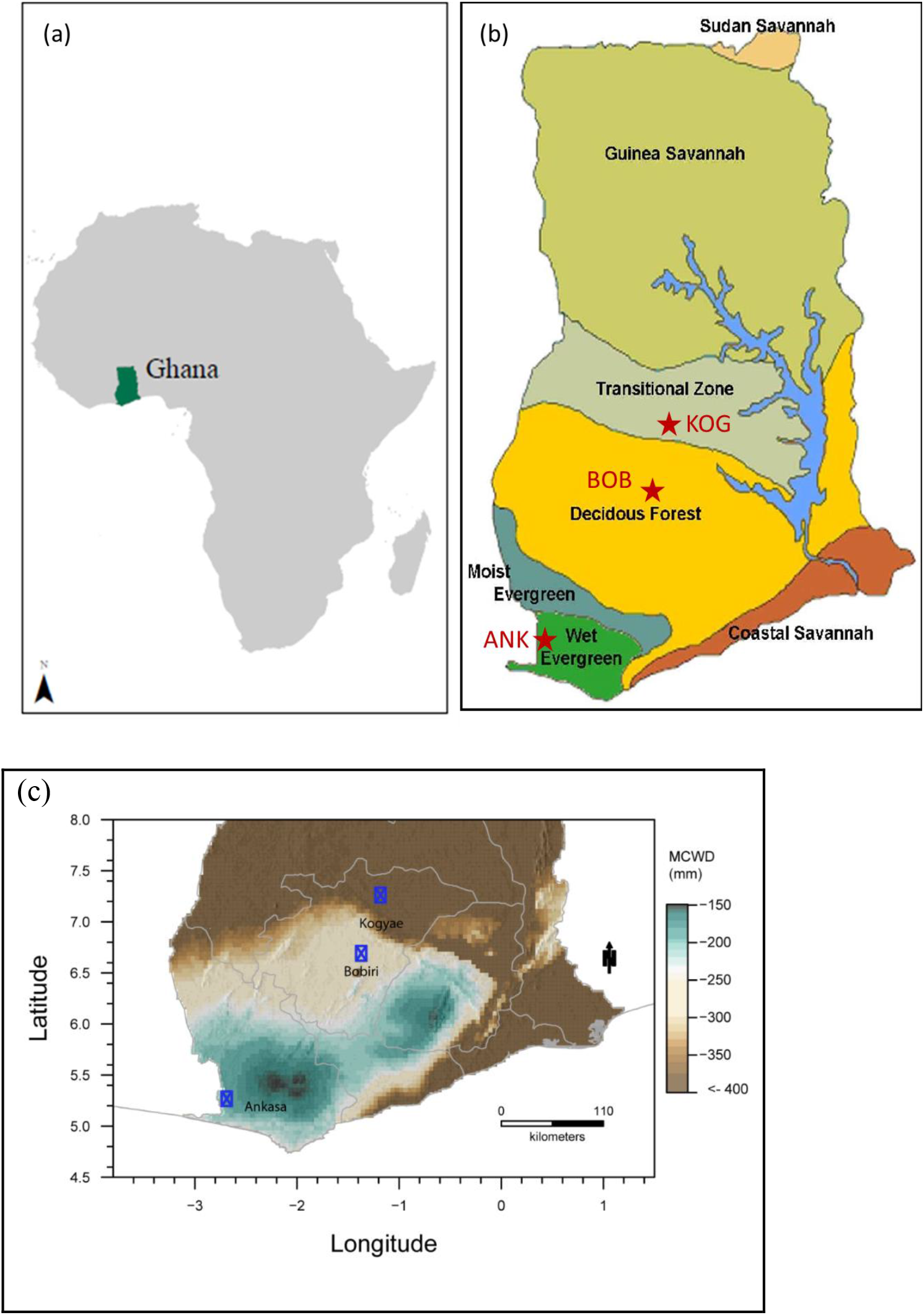
Map of the location of (a) Ghana within the African continent (b) and the study sites and forest types in Ghana (Appiah et al., 2014). (c) showed study sites over a map of maximum climate water deficit (MCWD) (Aguirre-Gutiérrez et al., 2019).

**Figure S 2.**
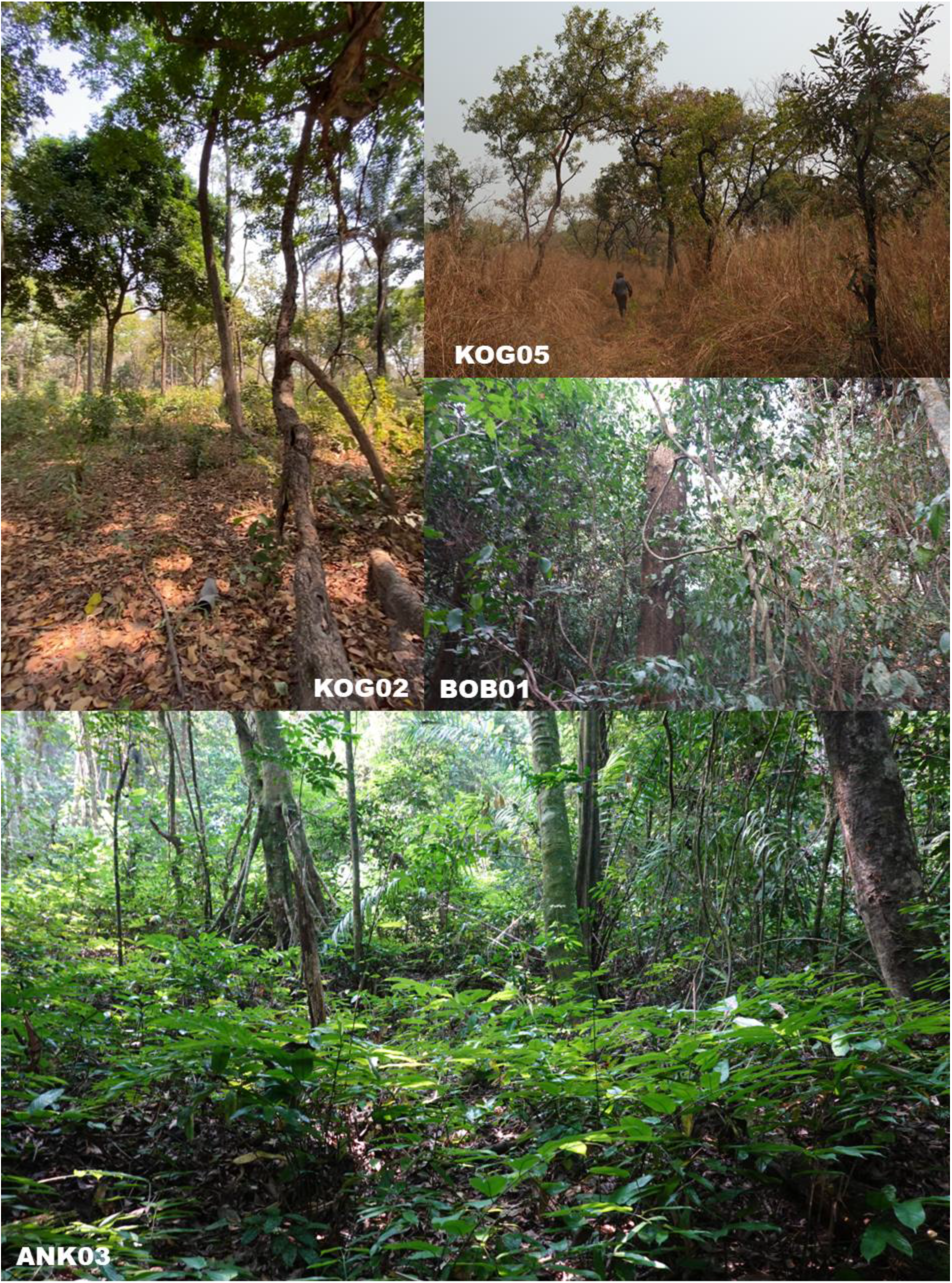
Photo of KOG02 (dry site), KOG05(dry site), ANK03 (wet site), and BOB01 (mid site). All photos were taken in January 2022 by Huanyuan Zhang-Zheng

**Table S 1.**
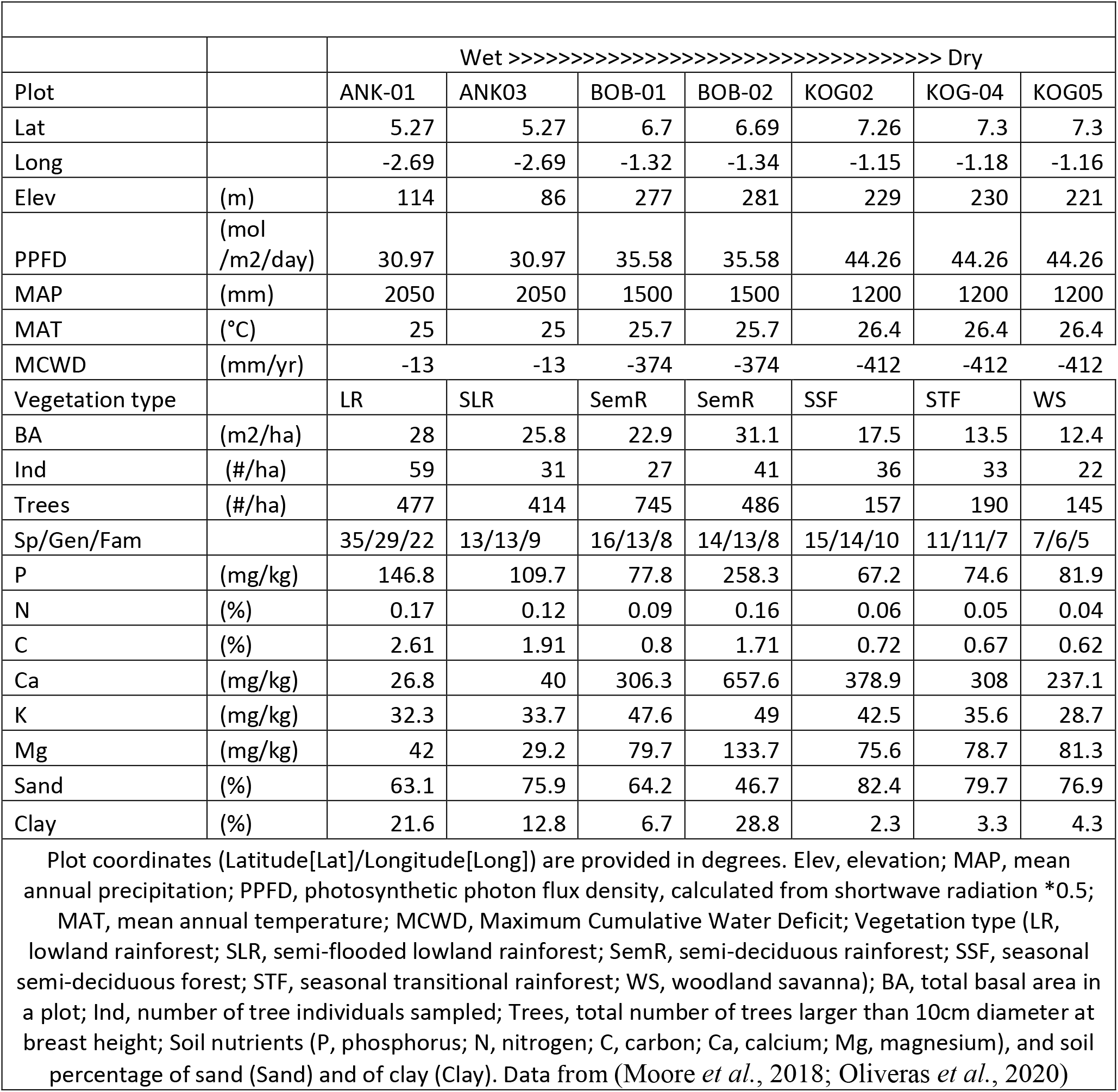
Summary information of plot characteristics

**Table S 2.**
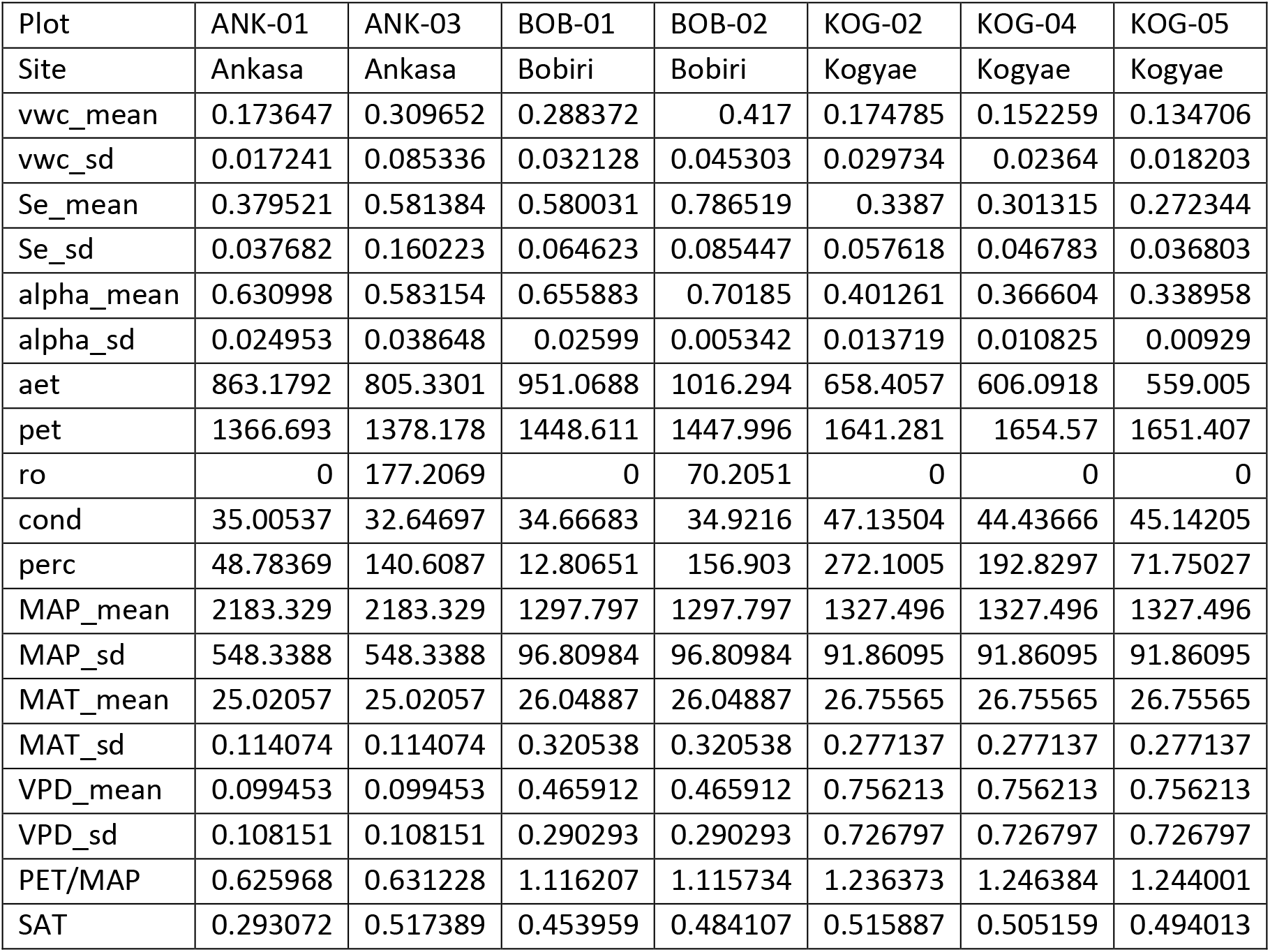
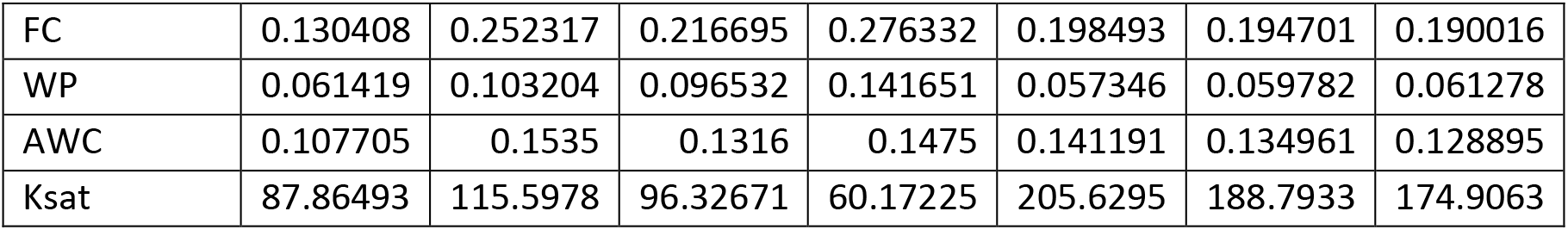
Rsplash model outputs, which simulate hydrology from climate, typography and soil property. vwc_mean and vwc_sd are mean and standard deviation of soil volumetric water content (unitless fraction); Se_mean and Se_sd are the mean and standard deviation of relative soil moisture saturation (Θ) (unitless fraction); Alpha_mean and Alpha_sd are mean and standard deviation of vegetation water stress index (α) (unitless fraction, calculated as AET/PET); Aet is actual evapotranspiration (mm year-1); Pet is potential evapotranspiration (mm year-1); ro is runoff (mm year-1); cond is condensation (mm year-1); MAP is mean annual precipitation (mm year-1), MAT is mean annual air temperature (degree Celsius), VPD is vapor pressure deficit (kpa, annual stats); SAT: volumetric water content at saturation m3 m-3; perc: percolation or deep drainage, or vertical drainage (mm/year); FC is volumetric water content at 33kPa (field capacity) m^3/m^3; WP, volumetric water content at 1500kPa (permanent wilting point) m^3/m^3; AWC: plant available volumetric water content (FC-WP); Ksat, Saturated hydraulic conductivity (mm/hr)

## Appendix 3 Information associated with leaf economy

Leaf economy traits have been published (Gvozdevaite *et al*., 2018; Oliveras *et al*., 2020). Data reported in this study are slightly different to the above two studys because there are more sampling in this study. Samples were taken each month from October 2014 to September 2016. Data here are provided for completeness and for future researchers ‘convenience.

**Figure S 3.**
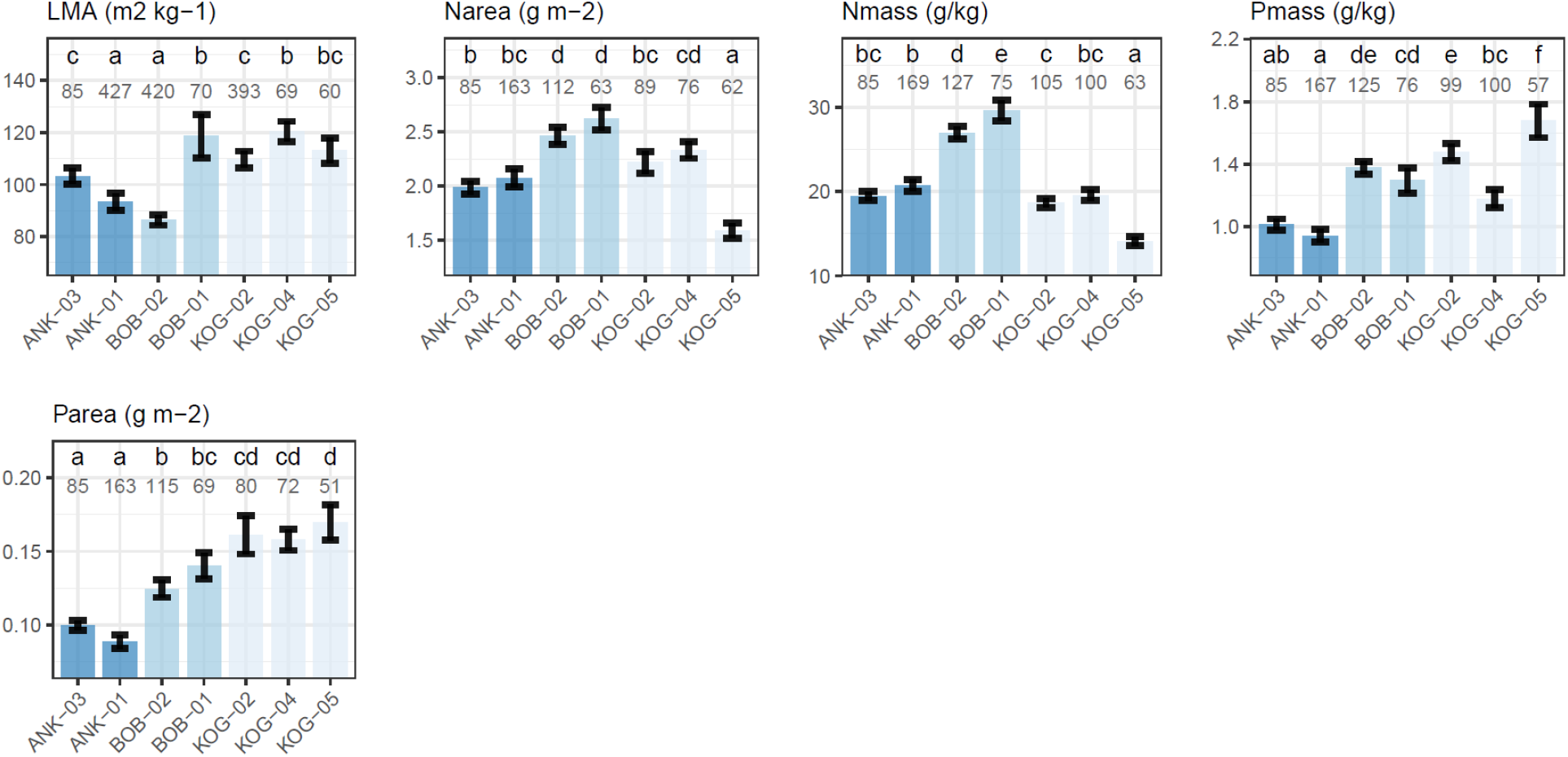
Community weighted mean (with standard error) of variables associated with leaf economy from wet to dry plots. Plots were ordered from left to right according to the description in the first paragraph of results. The number denotes the number of samples, which could be a leaf, a branch or a tree etc. The letters denote significance (P<0.05) in plot-to-plot difference. (Oliveras et al., 2020)

## Appendix 4 Report on Hypothesis 15

We hypothesized that the product of K_P_ (specific xylem hydraulic conductivity) and A_S_/A_L_ (sapwood to leaf area) vary less than K_P_ or A_S_/A_L_ themselves, and there is a trade-off (negative correlation) between K_P_ and AS/AL. As the trade-off between Ks (well associated with K_P_) and A_S_/A_L_ has been observed on a global scale (Mencuccini *et al*., 2019b), here we also plot K_P_ versus A_S_/A_L_ for readers’ convenience in comparison with measurements from Ghana aridity gradient. We estimated K_P_ for species reported in (Mencuccini *et al*., 2019b) by collecting vessel diameter and vessel density from XFT database (Choat *et al*., 2012), with the same calculation method as K_P_ of Ghana aridity gradient.

For Ghana, both hypotheses were rejected, as we see a positive correlation between K_P_ and A_S_/A_L_ (slope = 0.95, R-squared: 0.0598, P-value: 0.0224) and the coefficient of variance is found largest for K_P_ * A_S_/A_L_.

For a global dataset (Mencuccini et al., 2019), there is a negative correlation between K_P_ and A_S_/A_L_ (slope =-0.638, R-squared: 0.153, P-value: <0.001) which agreed with the hypothesis but the coefficient of variance of K_P_ * A_S_/A_L_ is still larger than that of either K_P_ or A_S_/A_L_.

Therefore, hypothesis 15 in Table 1 is rejected in this study. The negative correlation between K_P_ and A_S_/A_L_ emerge on global scale probably because of confounding effect with other environmental variables. The different patterns emerged at different scale could also result from a Simpson’s paradox. For example, the drier sites (KOG) have higher K_P_, higher twig density and higher wood density than the wetter sites on site scale (Figure 2), but we also found K_P_ negatively correlated with twig density on species scale (Figure S5)

**Figure S 4.**
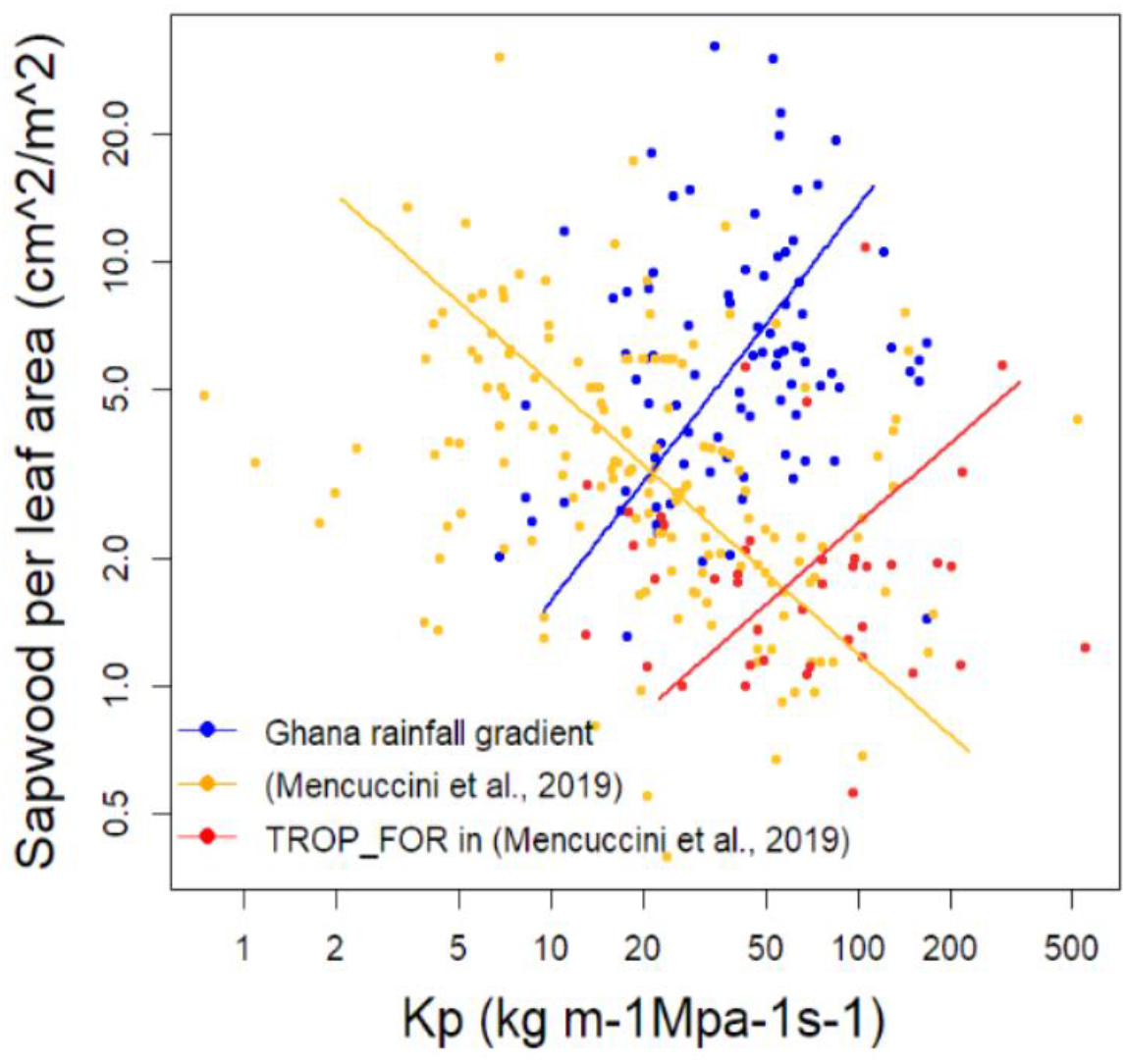
The correlation between sapwood to leaf area (AS/AL) and potential sapwood hydraulic conductivity (Kp) for Ghana aridity gradient (ANK, BOB and KOG all together) and species included in (Mencuccini et al., 2019b). The figure was drawn on species scale (one scatter point is one species).

**Figure S 5.**
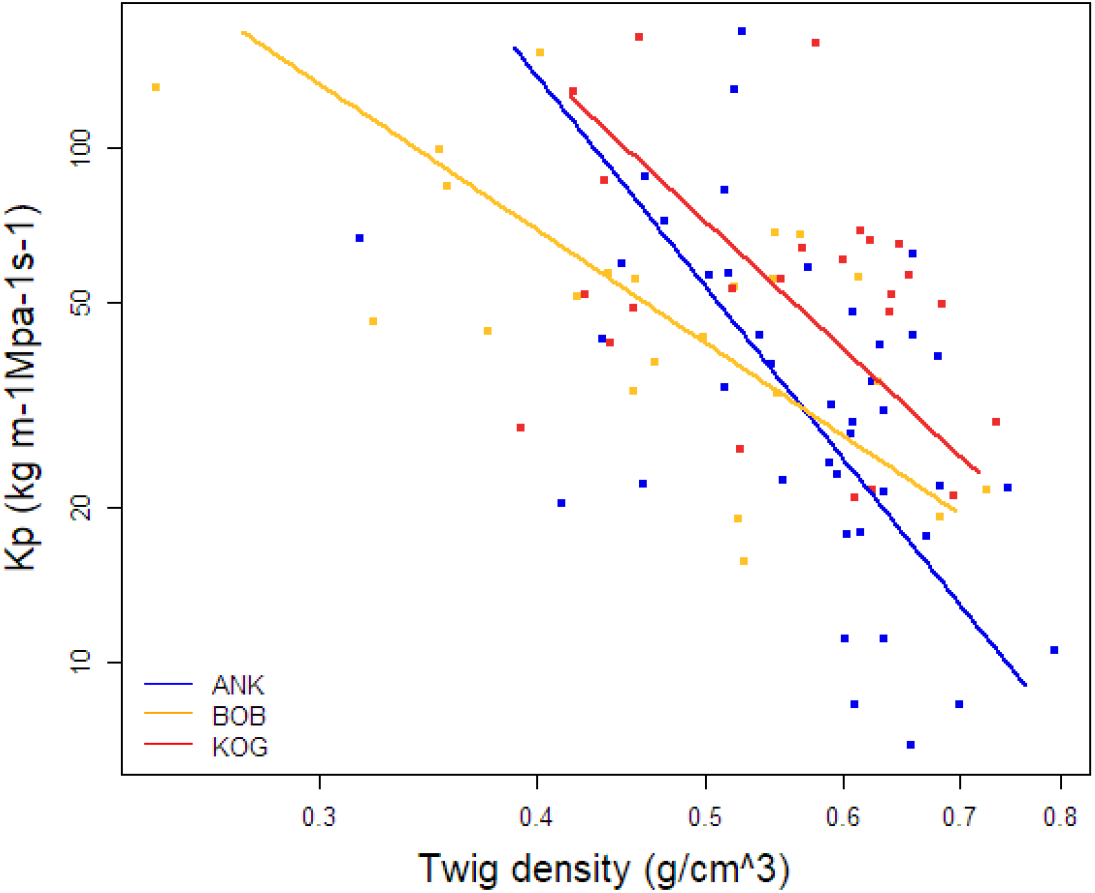
The correlation between twig density (g/cm3) and potential sapwood hydraulic conductivity (Kp) for site ANK, BOB and KOG. The figure was drawn on species scale (one scatter point is one species).

**Figure S 6.**
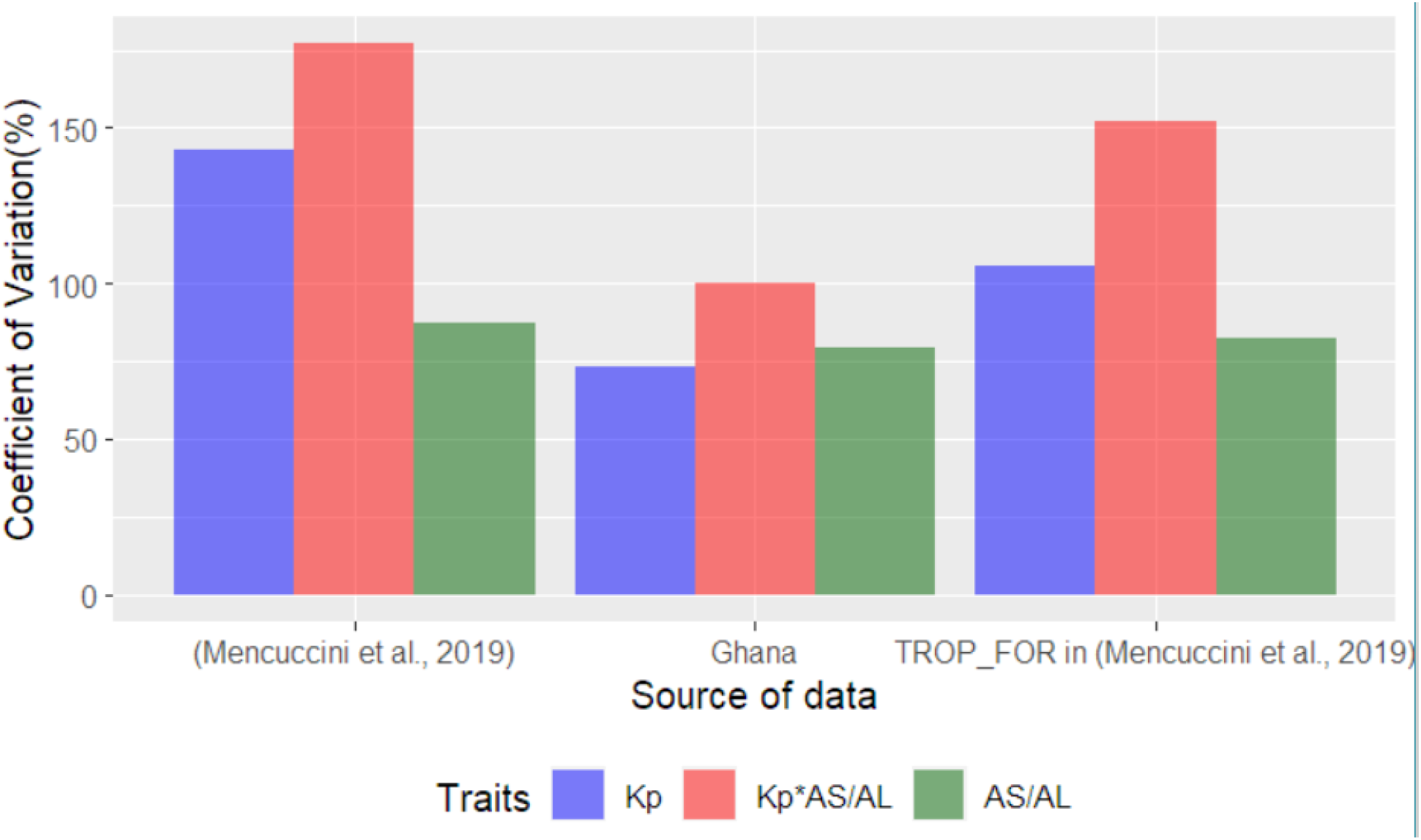
Coefficient of variation (%) for data points shown in figure S6, potential sapwood hydraulic conductivity (Kp), sapwood area to leaf area (AS/AL) and the product of Kp and AS/AL

**Figure S 7.**
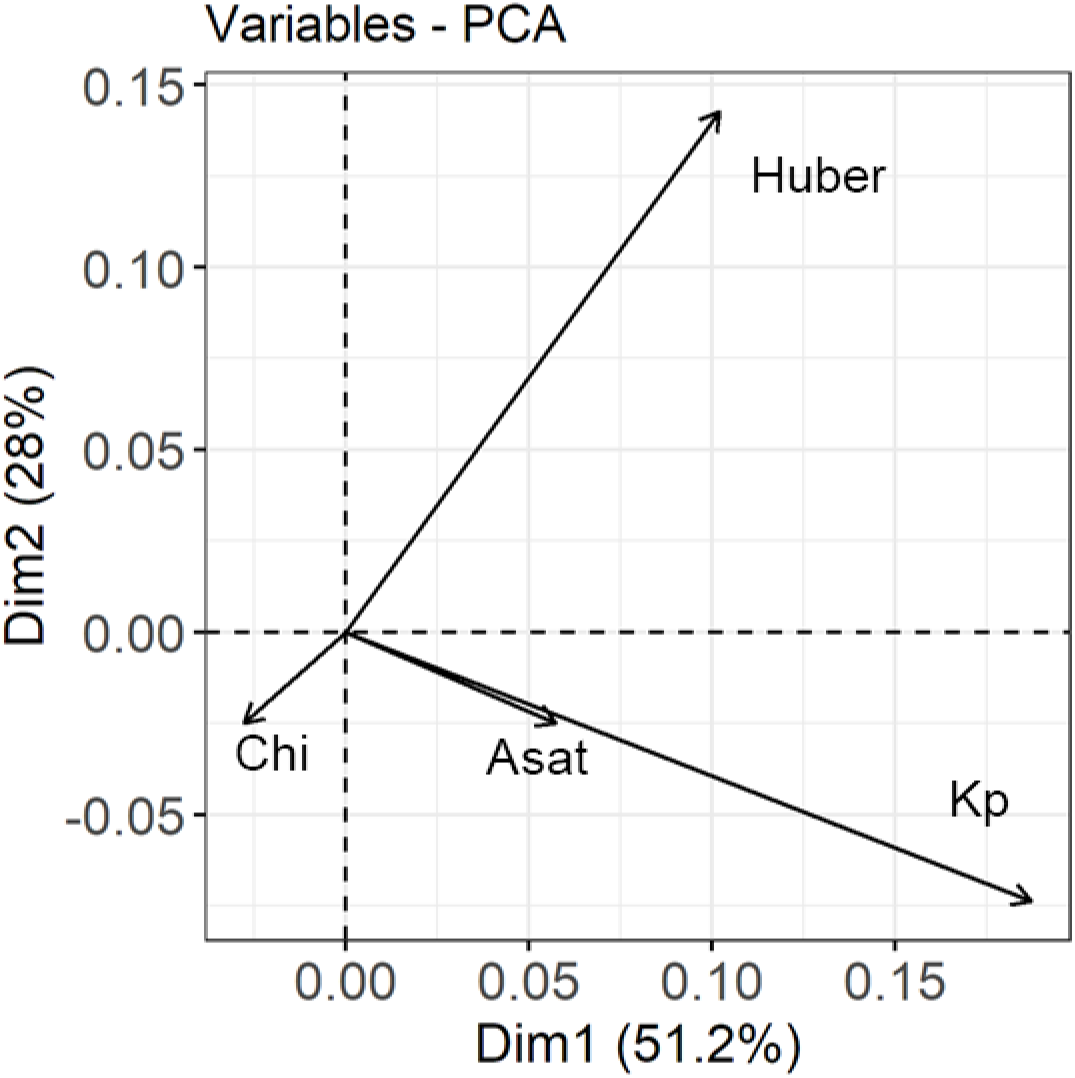
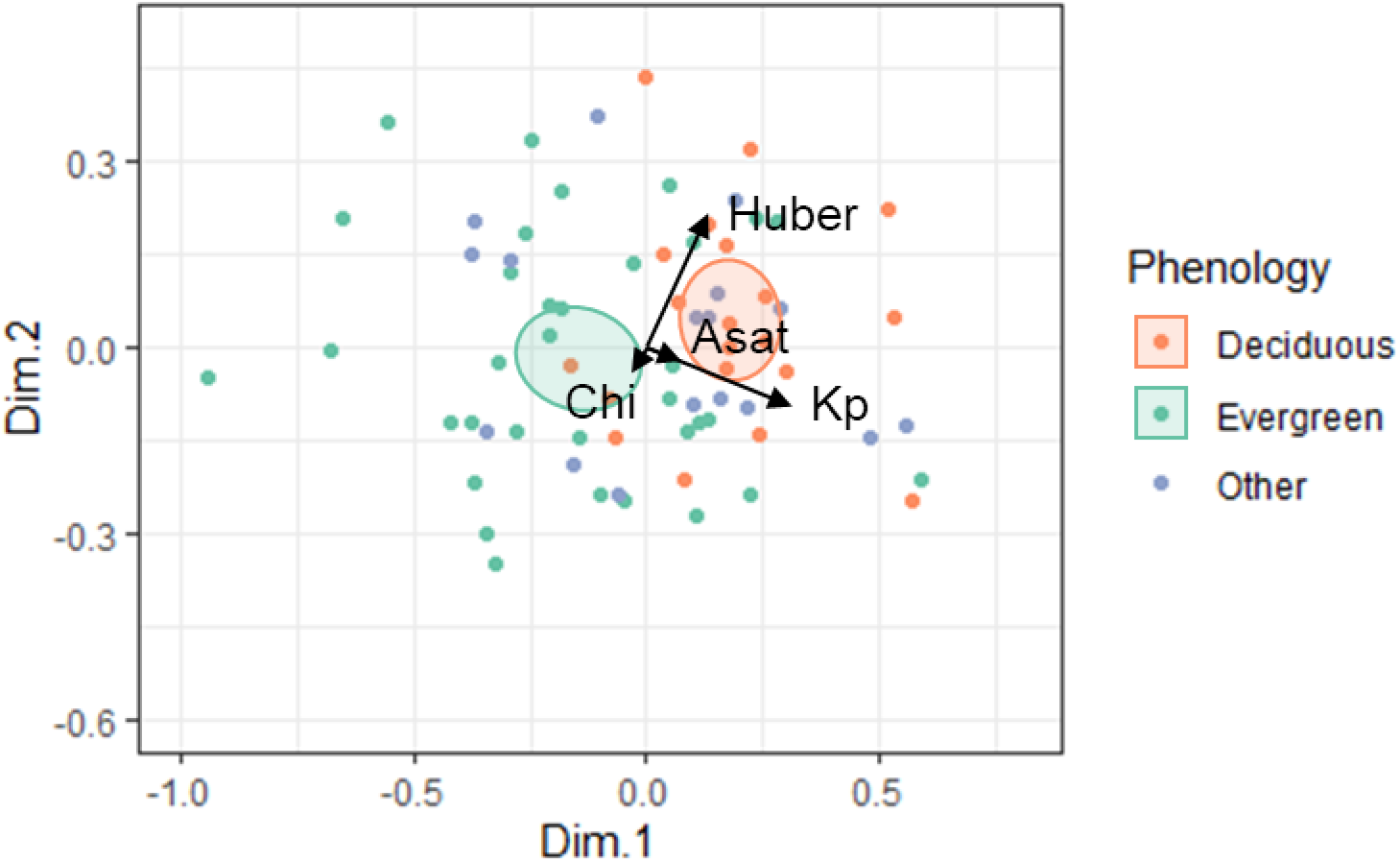
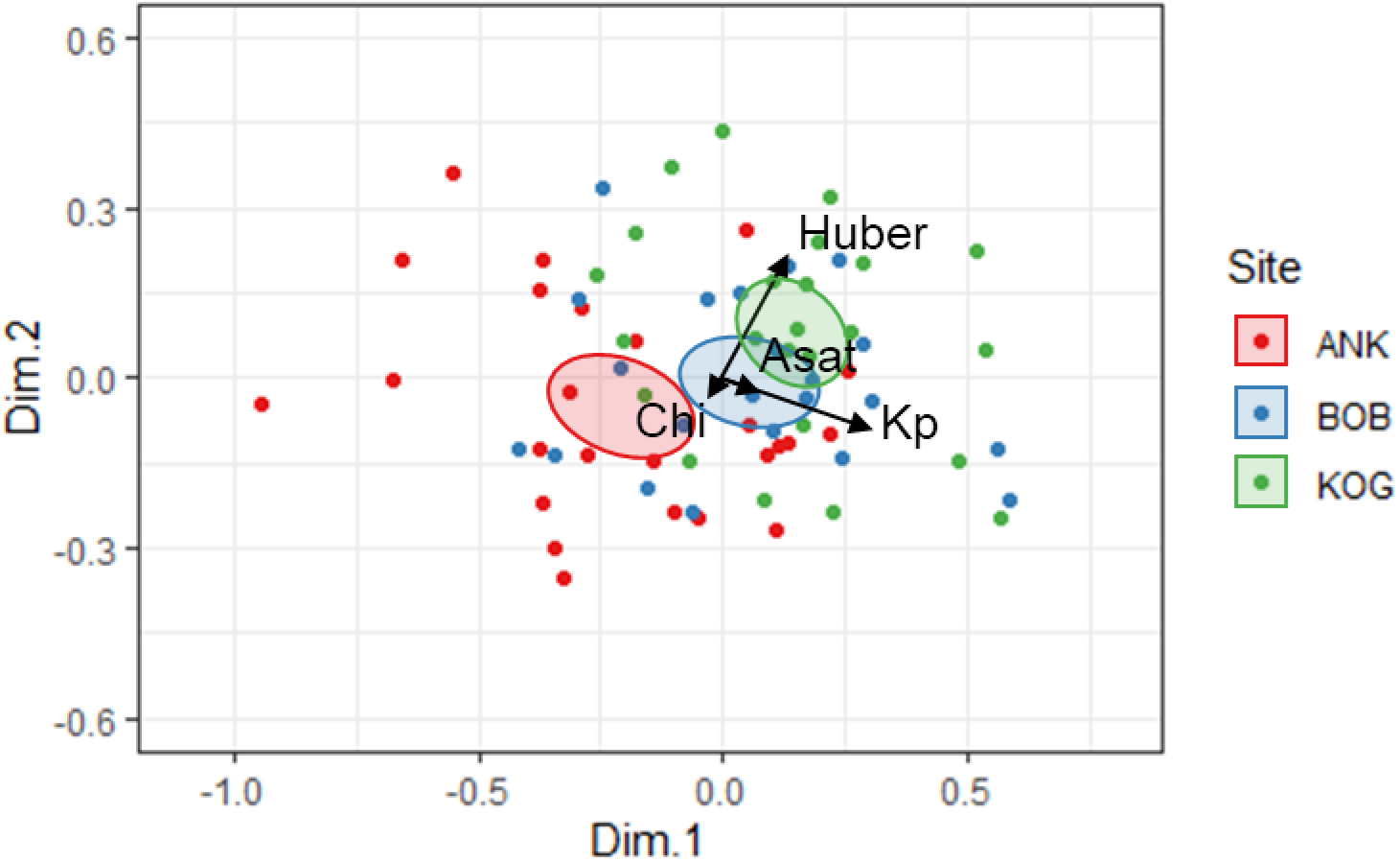
Principal components analysis for A_S_/A_L_, ci/ca, Asat and Kp. Pleaser also see Figure 2

**Figure S 8.**
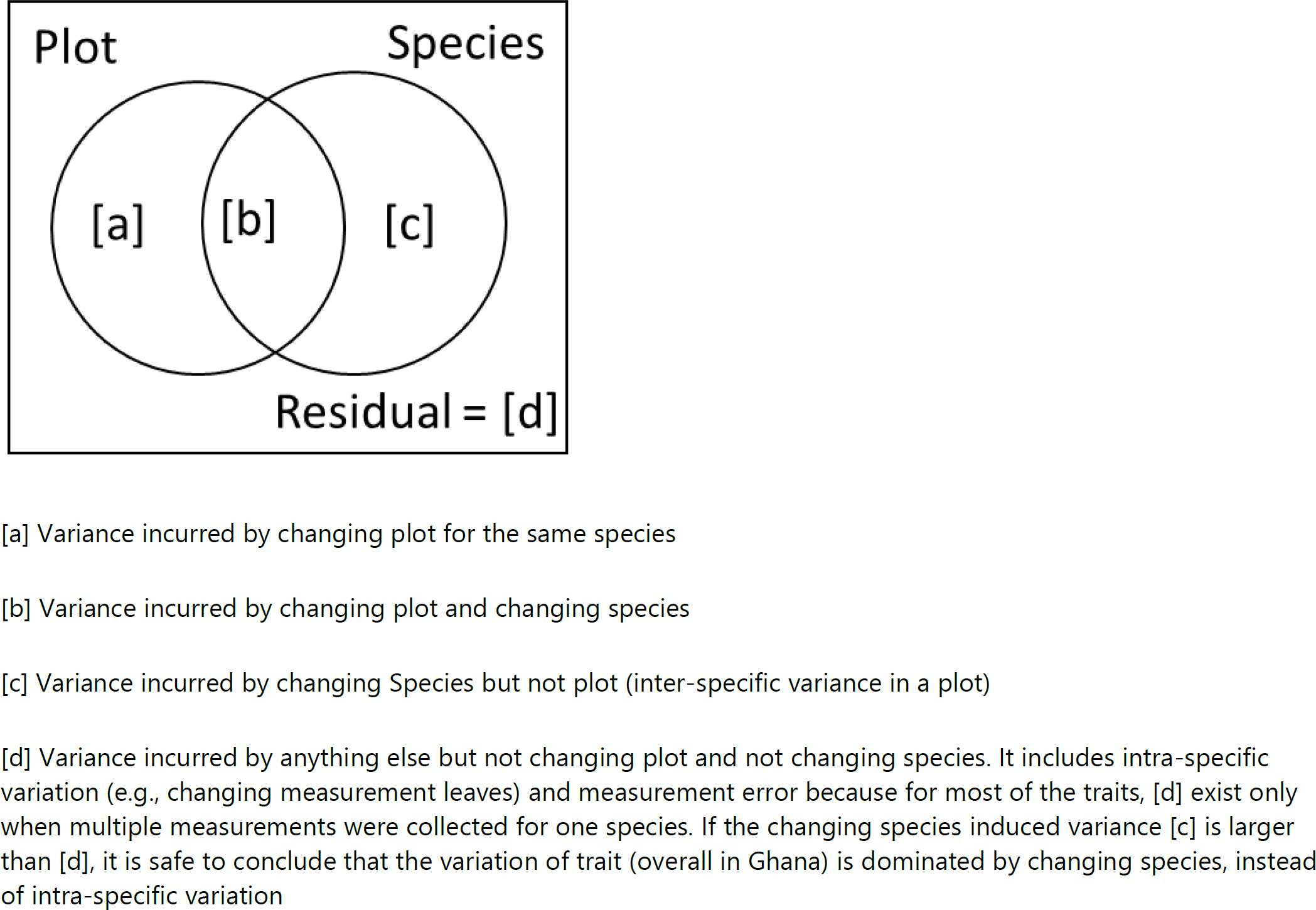
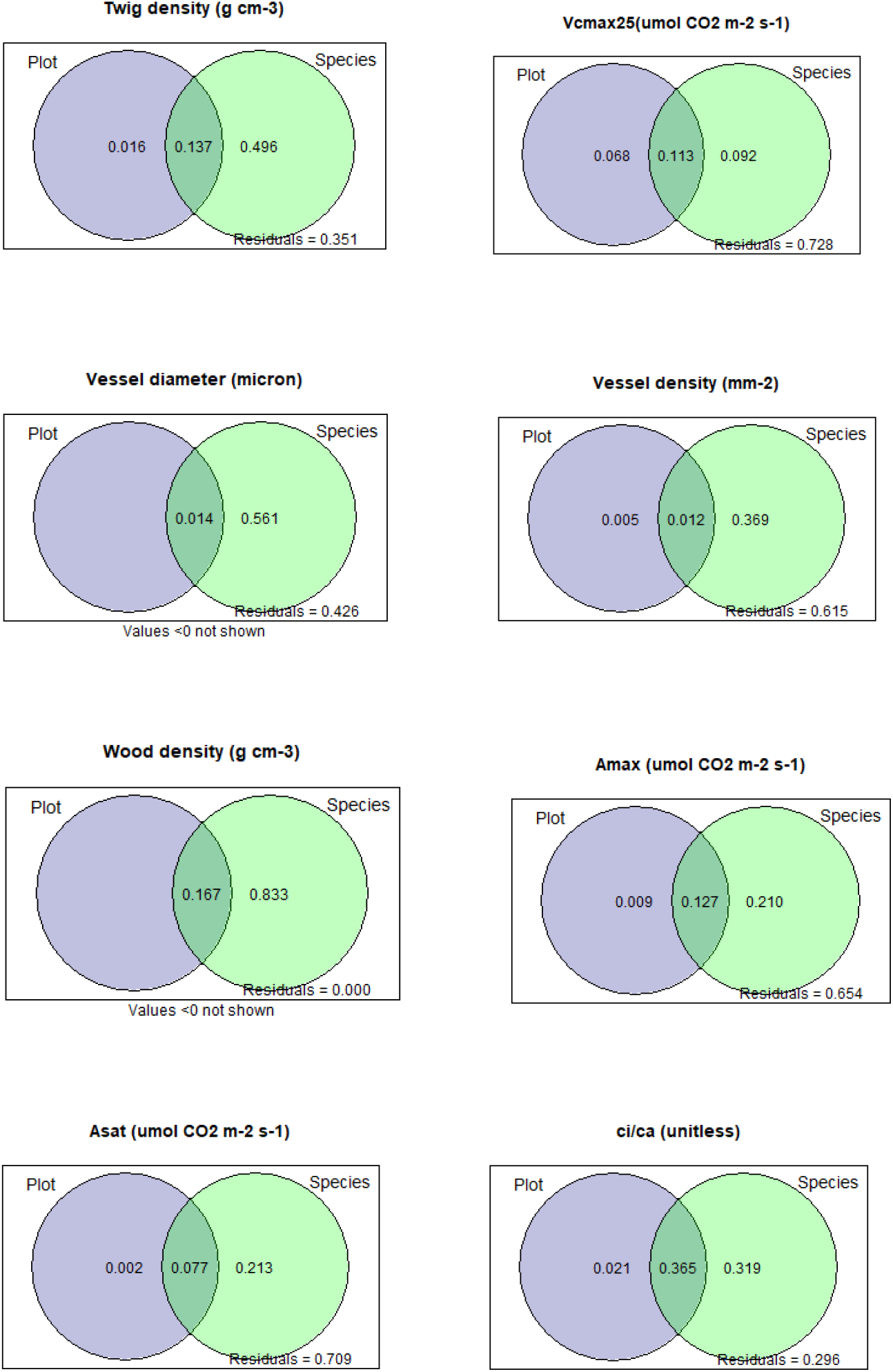
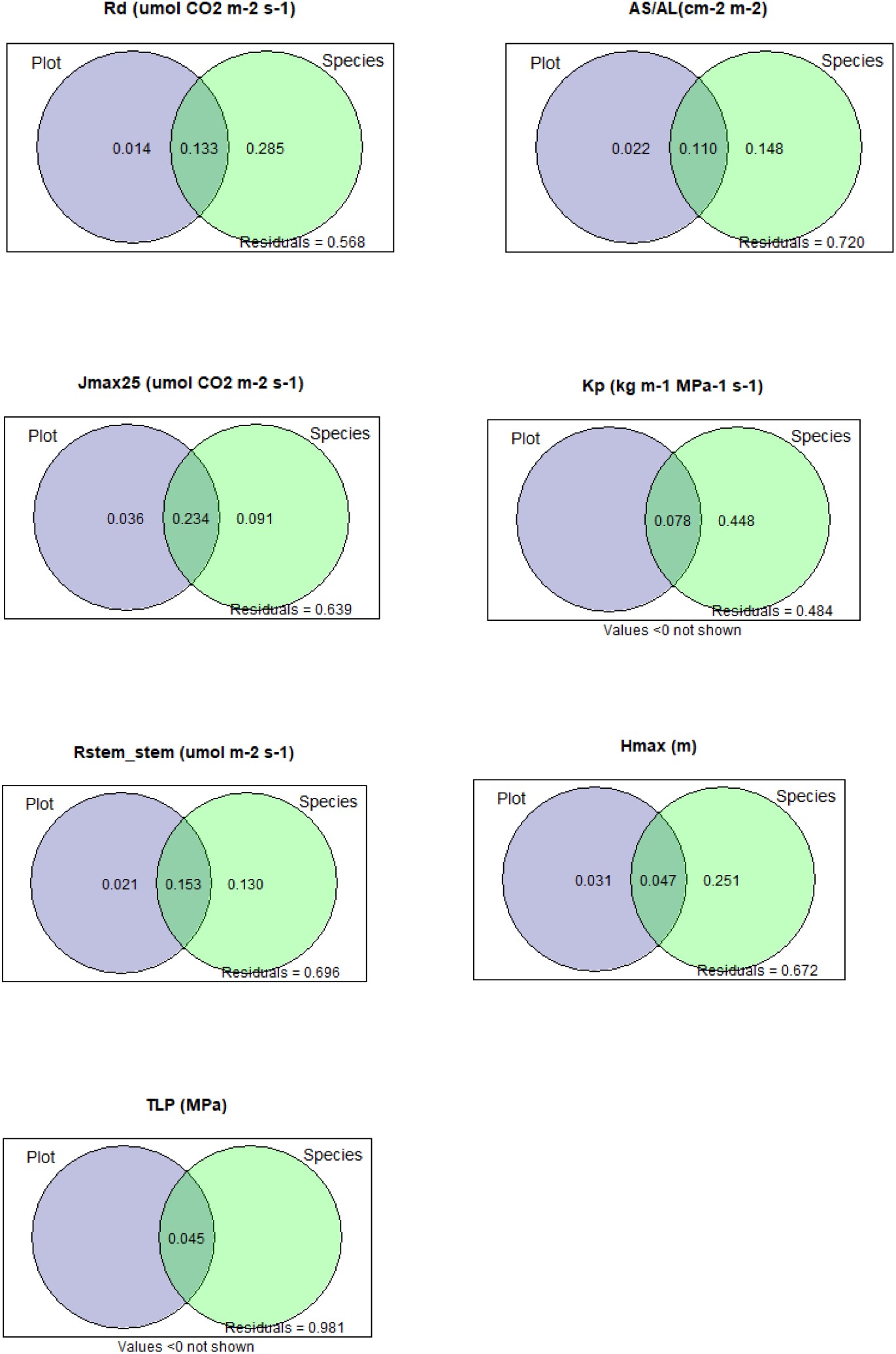
Variance partitioning into plot and species. Please see table 1 for definition of traits. Meanings of each number in the circles are explained in the top panel. Note that Vcmax, LMA Narea and Parea from the same plots were published in (Gvozdevaite et al., 2018), and Asat Amax LMA, Nmass and Pmass from the same plots were published in (Oliveras et al., 2020). Values are not mathematically identical due to (1) different methods of variance partitioning and (2) one more year sampling than the previous publications

## Appendix 5 Other figures

**Figure S 9.**
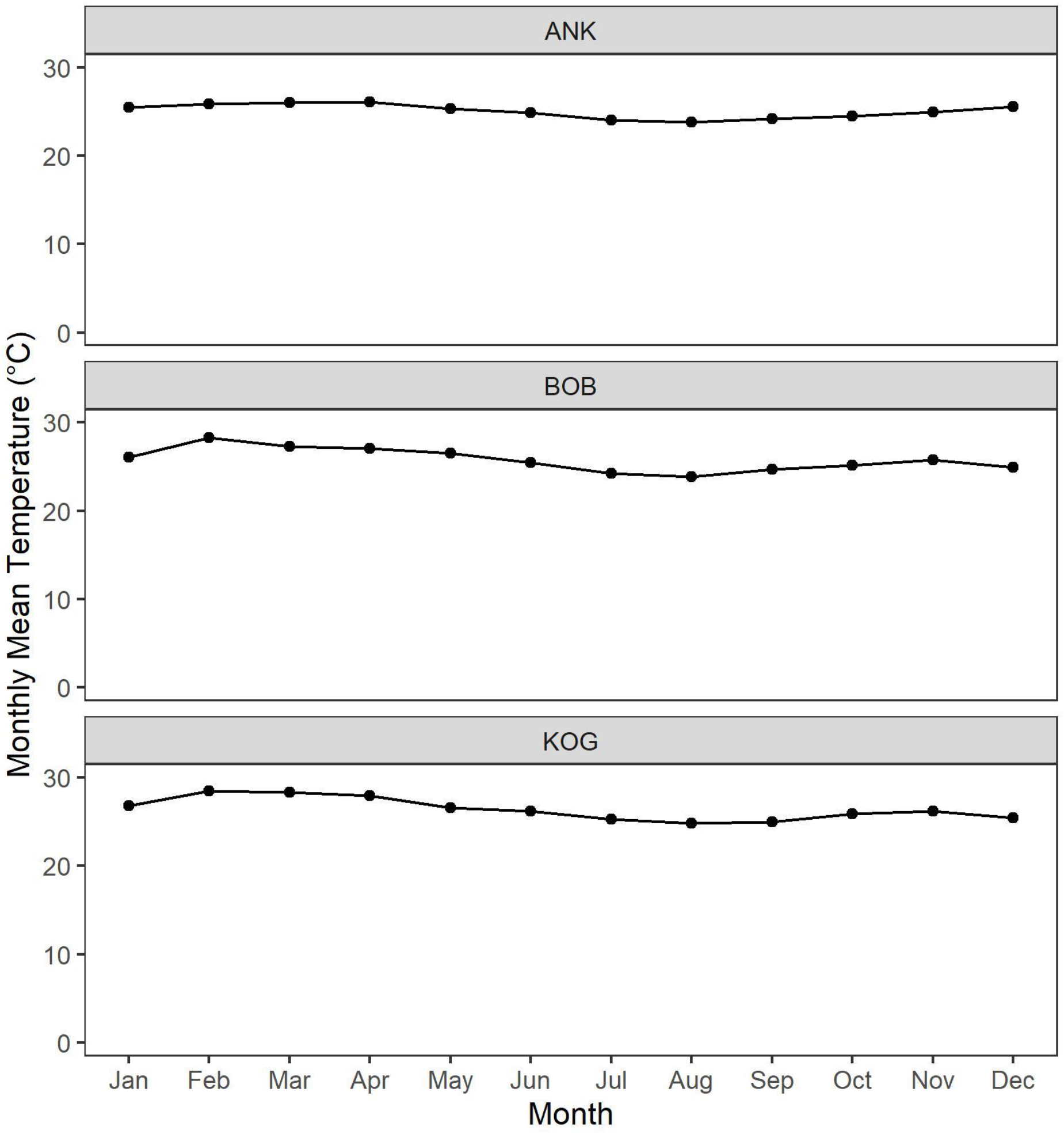
Monthly mean temperature at study sites ANK (wet site), BOB (mid site) and KOG (dry site), measured by in-situ climatological stations.

**Figure S 10.**
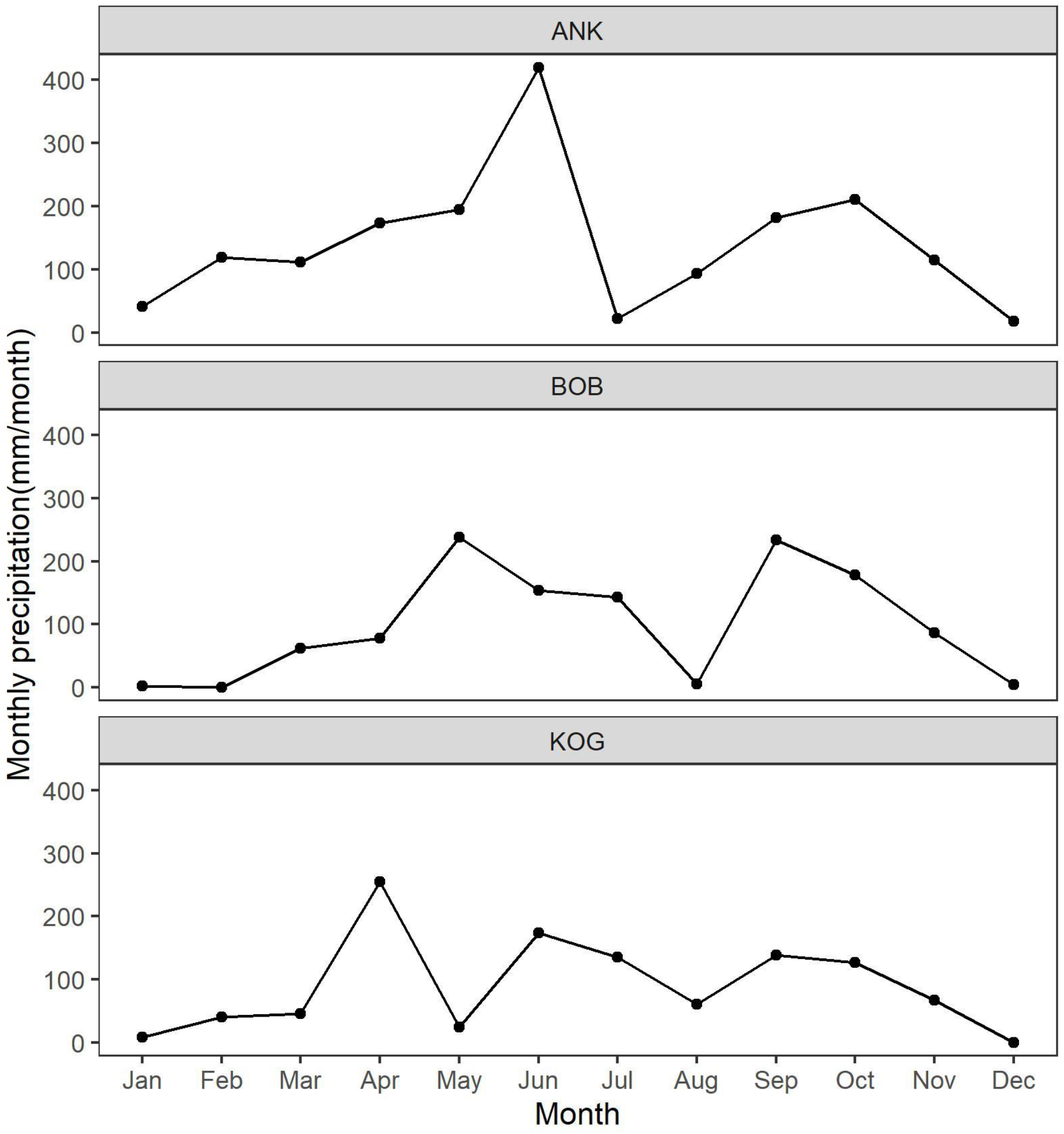
Monthly precipitation (mm/month) at study sites ANK (wet site, measured in 2011), BOB (mid site, measured in 2013) and KOG (dry site, measured in 2014), measured by in-situ climatological stations.

