## Supplementary for "Photosynthetic and water transport strategies of plants along a tropical forest aridity gradient: a test of optimality theory"

#### Appendix 1 Field sampling protocol

ci/ca (unitless) were estimated from leaf δ13C measurements (the stable isotope ratio relative to a standard material). We first estimated Δ13C the difference between the leaf stable isotope ratio and the atmospheric stable isotope ratio at that place and time according to (Cornwell *et al.*, 2016). Then we estimated ci/ca by equation 11 in (Peng *et al.*, 2020).

Soil volumetric water content (%) was measured in the field over the depth in the forest over the depth 0–12 cm.

For LMA (m-2 kg-1), Asat (umol $CO_2$ m-2 s-1) and Amax (umol $CO_2$ m-2 s-1), see supporting information in (Oliveras *et al.*, 2020).

For Parea (g m-2), Narea (g m-2), Nmass (g/kg), Pmass (g/kg), Rd (umol $CO_2$ m-2 s-1), $J_{max25}$ (umol $CO_2$ m-2 s-1) and $V_{cmax25}$ (umol $CO_2$ m-2 s-1), please see chapter 4 of (Gvozdevaite, 2018).

For AS/AL (cm-2 m-2), twig density (g cm-3), vessel density (mm-2), vessel diameter (micron) and Kp (kg m-1 MPa-1 s-1), Hmax (m) please see supplementary of (Aguirre-Gutiérrez *et al.*, 2019)

For TLP (MPa), see chapter four of (Raab, 2020)

Wood density was provided by Forestplot.net who sourced information from (Zanne *et al.*, 2009). In appendix 4, we also compared data collected from Ghana with AS/AL from (Mencuccini *et al.*, 2019)and vessel diameter from xylem functional traits database (Choat *et al.*, 2012).

Stem respiration per steam area (Rs_stem) was measured using a closed dynamic chamber method, from 25 trees distributed evenly throughout each plot at 1.3 m height with an IRGA (EGM-4) and soil respiration chamber (SRC-1) connected to a permanent collar, sealed to the tree bole surface. On plot scale, Rs_stem was multiplied by AS/AL to calculate stem respiration per leaf area (Rs_leaf).

#### Appendix 2 Study Sites, the aridity gradient

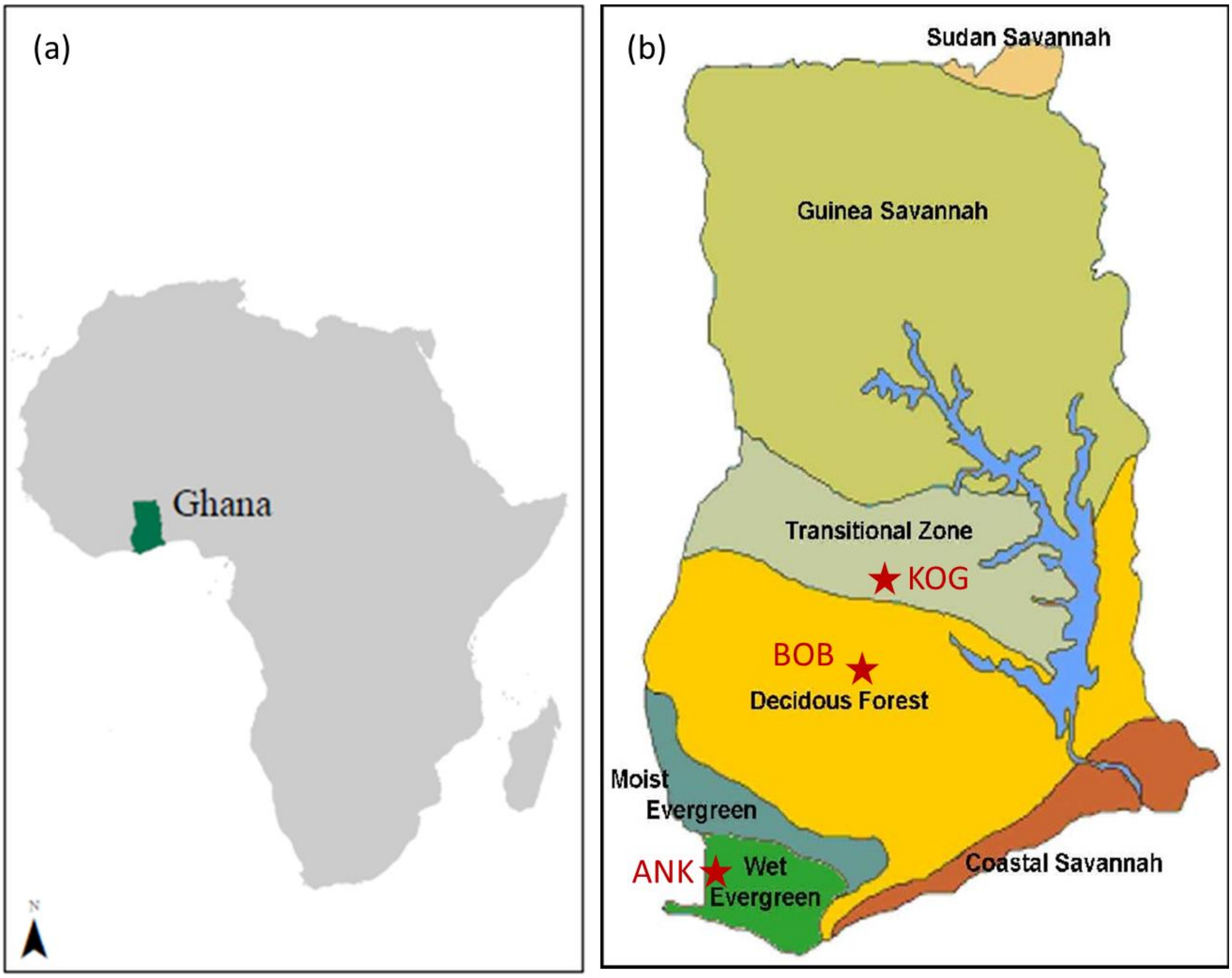


*Figure S 1 Map of the location of (a) Ghana within the African continent (b) and the study plots and forest types in Ghana* (Appiah *et al.*, 2014).

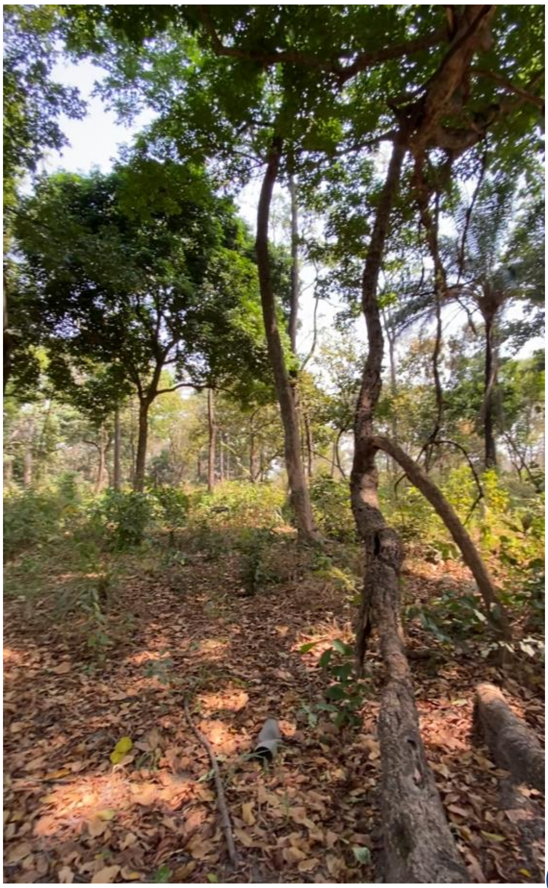
(KOG02)

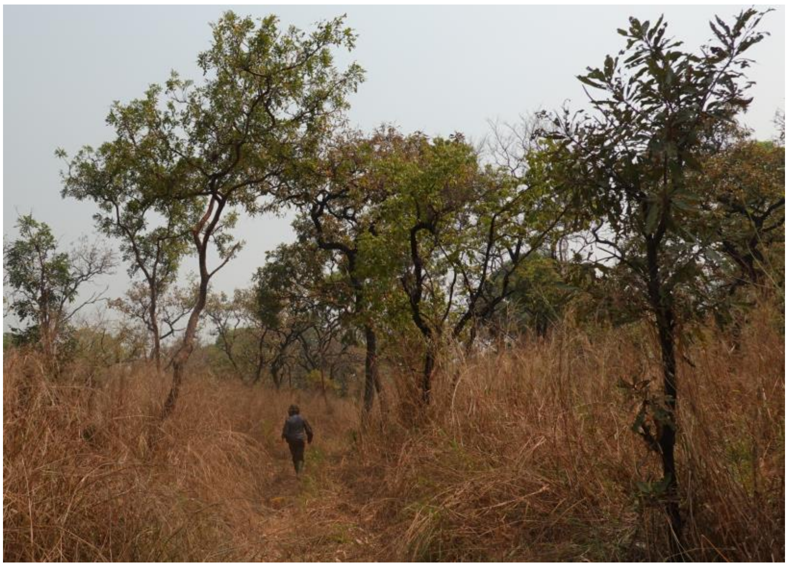

(KOG05)

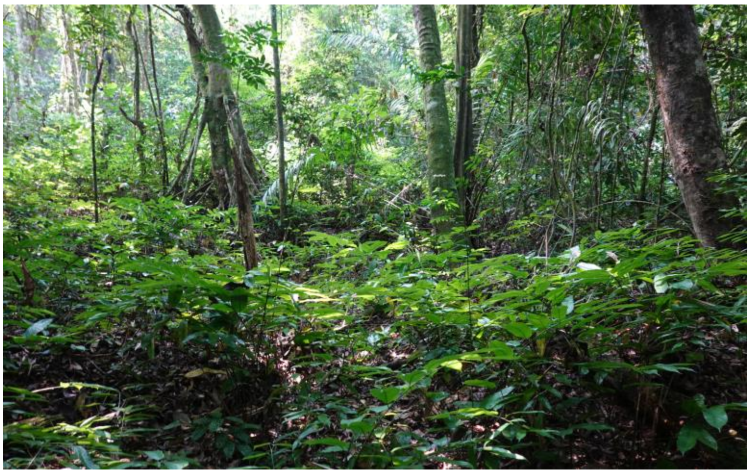

(ANK03)

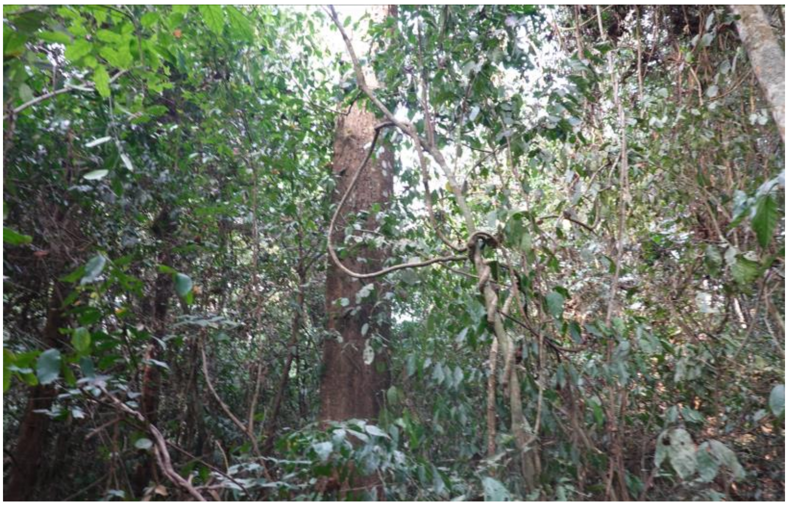

(BOB01)

*Figure S 2 Photo of KOG02 (dry site), KOG05(dry site), ANK03 (wet site), and BOB01 (mid site). All photos taken in January 2022 by Huanyuan Zhang-Zheng*

*Table S 1 Summary information of plot characteristics*

| | | | | | | | | |
|---|---|---|---|---|---|---|---|---|
| | | Wet >>>>>>>>>>>>>>>>>>>>>>>>>>>>>>>>>>> Dry | | | | | | |
| Plot | | ANK-01 | ANK03 | BOB-01 | BOB-02 | KOG02 | KOG-04 | KOG05 |
| Lat | | 5.27 | 5.27 | 6.7 | 6.69 | 7.26 | 7.3 | 7.3 |
| Long | | -2.69 | -2.69 | -1.32 | -1.34 | -1.15 | -1.18 | -1.16 |
| Elev | (m) | 114 | 86 | 277 | 281 | 229 | 230 | 221 |
| MAP | (mm) | 2050 | 2050 | 1500 | 1500 | 1200 | 1200 | 1200 |
| MAT | (°C) | 25 | 25 | 25.7 | 25.7 | 26.4 | 26.4 | 26.4 |
| MCWD | mm/year | -13 | -13 | -374 | -374 | -412 | -412 | -412 |
| Vegetation type | | LR | SLR | SemR | SemR | SSF | STF | WS |

| | | | | | | | | |
|---|---|---|---|---|---|---|---|---|
| BA | (m2/ha) | 28 | 25.8 | 22.9 | 31.1 | 17.5 | 13.5 | 12.4 |
| Ind | | 59 | 31 | 27 | 41 | 36 | 33 | 22 |
| Sp/Gen/Fam | | 35/29/22 | 13/13/9 | 16/13/8 | 14/13/8 | 15/14/10 | 11/11/7 | 7/6/5 |
| P | (mg/kg) | 146.8 | 109.7 | 77.8 | 258.3 | 67.2 | 74.6 | 81.9 |
| N | (%) | 0.17 | 0.12 | 0.09 | 0.16 | 0.06 | 0.05 | 0.04 |
| C | (%) | 2.61 | 1.91 | 0.8 | 1.71 | 0.72 | 0.67 | 0.62 |
| Ca | (mg/kg) | 26.8 | 40 | 306.3 | 657.6 | 378.9 | 308 | 237.1 |
| K | (mg/kg) | 32.3 | 33.7 | 47.6 | 49 | 42.5 | 35.6 | 28.7 |
| Mg | (mg/kg) | 42 | 29.2 | 79.7 | 133.7 | 75.6 | 78.7 | 81.3 |
| Sand | (%) | 63.1 | 75.9 | 64.2 | 46.7 | 82.4 | 79.7 | 76.9 |
| Clay | (%) | 21.6 | 12.8 | 6.7 | 28.8 | 2.3 | 3.3 | 4.3 |

Plot coordinates (Latitude[Lat]/Longitude[Long]) are provided in degrees. Elev, elevation; MAP, mean annual precipitation; MAT, mean annual temperature; MCWD, Maximum Cumulative Water Deficit; Vegetation type (LR, lowland rainforest; SLR, semi-flooded lowland rainforest; SemR, semi-deciduous rainforest; SSF, seasonal semi-deciduous forest; STF, seasonal transitional rainforest; WS, woodland savanna); BA, total basal area in a plot; Ind, number of tree individuals; Soil nutrients (P, phosphorus; N, nitrogen; C, carbon; Ca, calcium; Mg, magnesium), and soil percentage of sand (Sand) and of clay (Clay). Data from (Moore *et al.*, 2018; Oliveras *et al.*, 2020)

*Table S2 Rsplash model outputs, which simulate hydrology from climate, typography and soil property. v*wc_mean and vwc_sd are mean and standard deviation of soil volumetric water content (unitless fraction); Se_mean and Se_sd are the mean and standard deviation of relative soil moisture saturation (Θ) (unitless fraction); Alpha_mean and Alpha_sd are mean and standard deviation of vegetation water stress index (α) (unitless fraction); Aet is actual evapotranspiration (mm year-1); Pet is potential evapotranspiration (mm year-1); ro is runoff (mm year-1); cond is condensation (mm year-1); MAP is mean annual precipitation (mm year-1), MAT is mean annual air temperature (degree Celsius), VPD is vapor pressure deficit (kpa, annual stats); SAT: volumetric water content at saturation m3 m-3; perc: percolation or deep drainage, or vertical drainage (mm/year); FC is volumetric water content at 33kPa (field capacity) m^3/m^3; WP, volumetric water content at 1500kPa (permanent wilting point) m^3/m^3; AWC: plant available volumetric water content (FC-WP); Ksat, Saturated hydraulic conductivity (mm/hr)

| Plot | ANK-01 | ANK-03 | BOB-01 | BOB-02 | KOG-02 | KOG-04 | KOG-05 |
|---|---|---|---|---|---|---|---|
| Site | Ankasa | Ankasa | Bobiri | Bobiri | Kogyae | Kogyae | Kogyae |
| vwc_mean | 0.173647 | 0.309652 | 0.288372 | 0.417 | 0.174785 | 0.152259 | 0.134706 |

| | | | | | | | |
|---|---|---|---|---|---|---|---|
| vwc_sd | 0.017241 | 0.085336 | 0.032128 | 0.045303 | 0.029734 | 0.02364 | 0.018203 |
| Se_mean | 0.379521 | 0.581384 | 0.580031 | 0.786519 | 0.3387 | 0.301315 | 0.272344 |
| Se_sd | 0.037682 | 0.160223 | 0.064623 | 0.085447 | 0.057618 | 0.046783 | 0.036803 |
| alpha_mean | 0.630998 | 0.583154 | 0.655883 | 0.70185 | 0.401261 | 0.366604 | 0.338958 |
| alpha_sd | 0.024953 | 0.038648 | 0.02599 | 0.005342 | 0.013719 | 0.010825 | 0.00929 |
| aet | 863.1792 | 805.3301 | 951.0688 | 1016.294 | 658.4057 | 606.0918 | 559.005 |
| pet | 1366.693 | 1378.178 | 1448.611 | 1447.996 | 1641.281 | 1654.57 | 1651.407 |
| ro | 0 | 177.2069 | 0 | 70.2051 | 0 | 0 | 0 |
| cond | 35.00537 | 32.64697 | 34.66683 | 34.9216 | 47.13504 | 44.43666 | 45.14205 |
| perc | 48.78369 | 140.6087 | 12.80651 | 156.903 | 272.1005 | 192.8297 | 71.75027 |
| MAP_mean | 2183.329 | 2183.329 | 1297.797 | 1297.797 | 1327.496 | 1327.496 | 1327.496 |
| MAP_sd | 548.3388 | 548.3388 | 96.80984 | 96.80984 | 91.86095 | 91.86095 | 91.86095 |
| MAT_mean | 25.02057 | 25.02057 | 26.04887 | 26.04887 | 26.75565 | 26.75565 | 26.75565 |
| MAT_sd | 0.114074 | 0.114074 | 0.320538 | 0.320538 | 0.277137 | 0.277137 | 0.277137 |
| VPD_mean | 0.099453 | 0.099453 | 0.465912 | 0.465912 | 0.756213 | 0.756213 | 0.756213 |
| VPD_sd | 0.108151 | 0.108151 | 0.290293 | 0.290293 | 0.726797 | 0.726797 | 0.726797 |
| PET/MAP | 0.625968 | 0.631228 | 1.116207 | 1.115734 | 1.236373 | 1.246384 | 1.244001 |
| SAT | 0.293072 | 0.517389 | 0.453959 | 0.484107 | 0.515887 | 0.505159 | 0.494013 |

| | | | | | | | |
|---|---|---|---|---|---|---|---|
| FC | 0.130408 | 0.252317 | 0.216695 | 0.276332 | 0.198493 | 0.194701 | 0.190016 |
| WP | 0.061419 | 0.103204 | 0.096532 | 0.141651 | 0.057346 | 0.059782 | 0.061278 |
| AWC | 0.107705 | 0.1535 | 0.1316 | 0.1475 | 0.141191 | 0.134961 | 0.128895 |
| Ksat | 87.86493 | 115.5978 | 96.32671 | 60.17225 | 205.6295 | 188.7933 | 174.9063 |

#### Appendix 3 Information associated with leaf economy

It has been shown that assimilation rate and $V_{cmax25}$ are positively correlated with leaf nitrogen by area (Narea) and by dry mass (Nmass) (Wright *et al.*, 2004; Niinemets *et al.*, 2009; Walker *et al.*, 2014). On a site scale, Narea and Nmass tend to be higher in drier and brighter sites (Dong *et al.*, 2017, 2020). All the above suggests that Nmass should increase toward drier sites along the aridity gradient. Leaf phosphorus by dry mass (Pmass) also tends to be higher in drier sites (Maire *et al.*, 2015; Gvozdevaite *et al.*, 2018), and. it has been predicted that LMA should increase with light and aridity (Wright *et al.*, 2005; Poorter *et al.*, 2010).

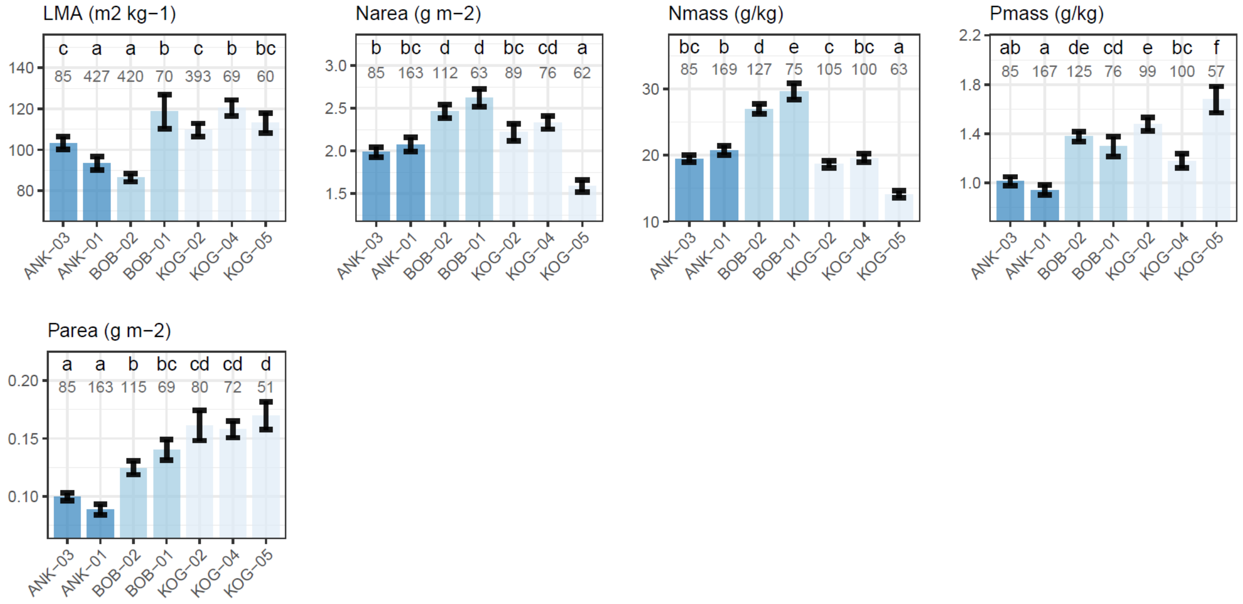


*Figure S3 Community weighted mean (with standard error) of variables associated with leaf economy from wet to dry plots. Plots were ordered from left to right according to the description in the first paragraph of results. The number denotes the number of samples, which could be a leaf, a branch or a tree etc. The letters denote significance (P<0.05) in plot-to-plot difference.*

### Appendix 4 Report on hypothesis 15

We hypothesized that the product of Kp (specific xylem hydraulic conductivity) and AS/AL (sapwood to leaf area) vary less than Kp or AS/AL themselves, and there is a trade-off (negative correlation) between Kp and AS/AL. As the trade-off between Ks (well associated with Kp) and AS/AL has been observed on a global scale (Mencuccini *et al.*, 2019), here we also plot Kp versus AS/AL for readers' convenience in comparison with measurements from Ghana aridity gradient. We estimated Kp for species reported in (Mencuccini *et al.*, 2019) by collecting vessel diameter and vessel density from XFT database (Choat *et al.*, 2012), with the same calculation method as Kp of Ghana aridity gradient.

For Ghana, both hypotheses were rejected, as we see a positive correlation between Kp and AS/AL (slope = 0.95, R-squared : 0.0598, P-value : 0.0224) and the coefficient of variance is found largest for Kp * AS/AL.

For a global dataset (Mencuccini et al., 2019), there is a negative correlation between Kp and AS/AL (slope =-0.638, R-squared: 0.153, P-value : <0.001) which agreed with the hypothesis but the coefficient of variance of Kp * AS/AL is still larger than that of either Kp or AS/AL.

Therefore, hypothesis 15 in Table 1 is rejected in this study. The negative correlation between Kp and AS/AL emerge on global scale probably because of confounding effect with other environmental variables. The different patterns emerged at different scale could also result from a Simpson's paradox. For example, the drier sites (KOG) have higher Kp, higher twig density and higher wood density than the wetter sites on site scale (Figure 2), but we also found Kp negatively correlated with twig density on species scale (Figure S5)

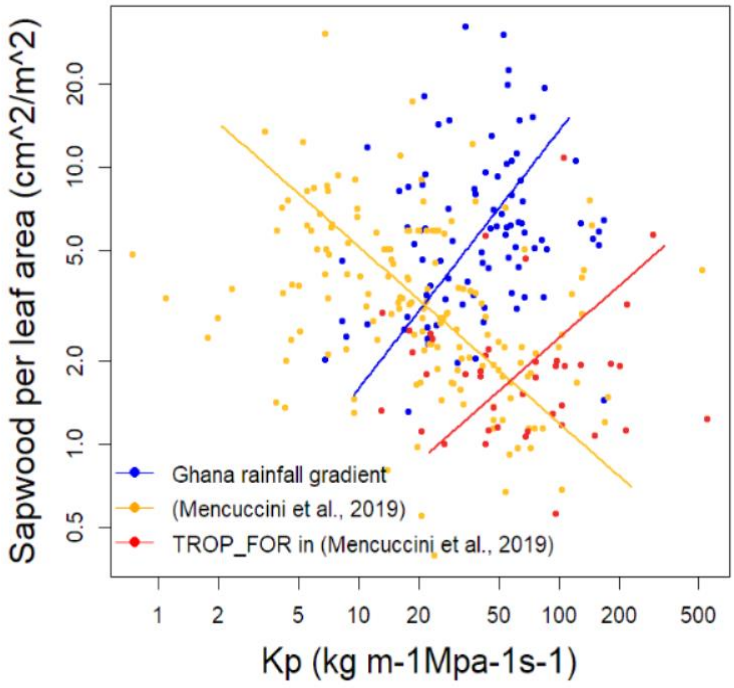


*Figure S4 The correlation between sapwood to leaf area (AS/AL) and potential sapwood hydraulic conductivity (Kp) for Ghana aridity gradient (ANK, BOB and KOG all together) and*

*species included in* (Mencuccini *et al.*, 2019)*. The figure was drawn on species scale (one scatter point is one species).*

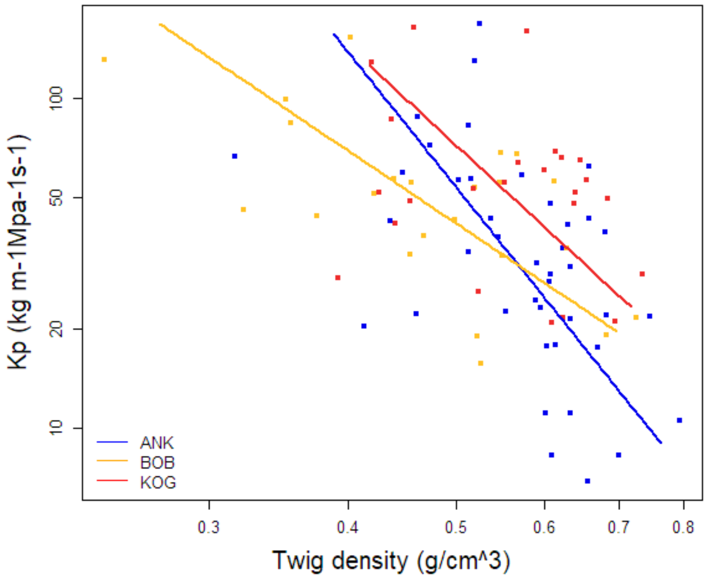


*Figure S5 The correlation between twig density (g/cm3) and potential sapwood hydraulic conductivity (Kp) for site ANK, BOB and KOG. The figure was drawn on species scale (one scatter point is one species).*

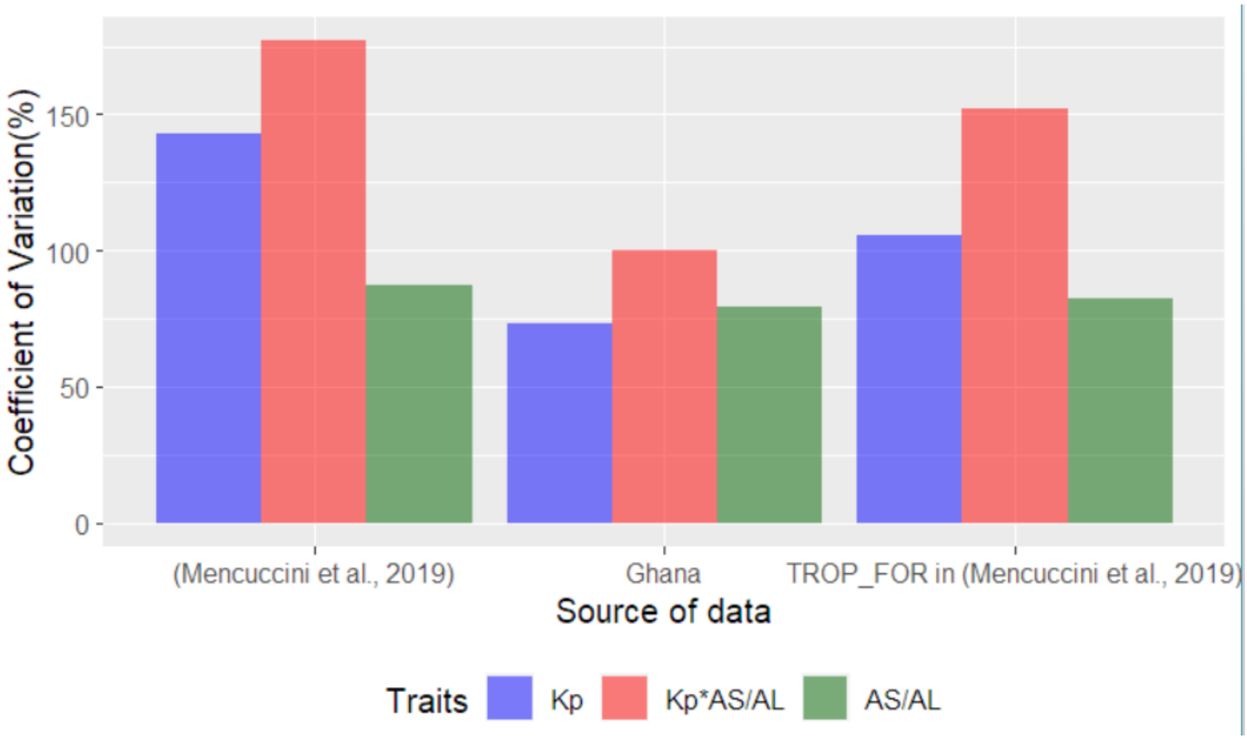


*Figure S 6 Coefficient of variation (%) for data points shown in figure S6, potential sapwood hydraulic conductivity (Kp), sapwood area to leaf area (AS/AL) and the product of Kp and AS/AL*

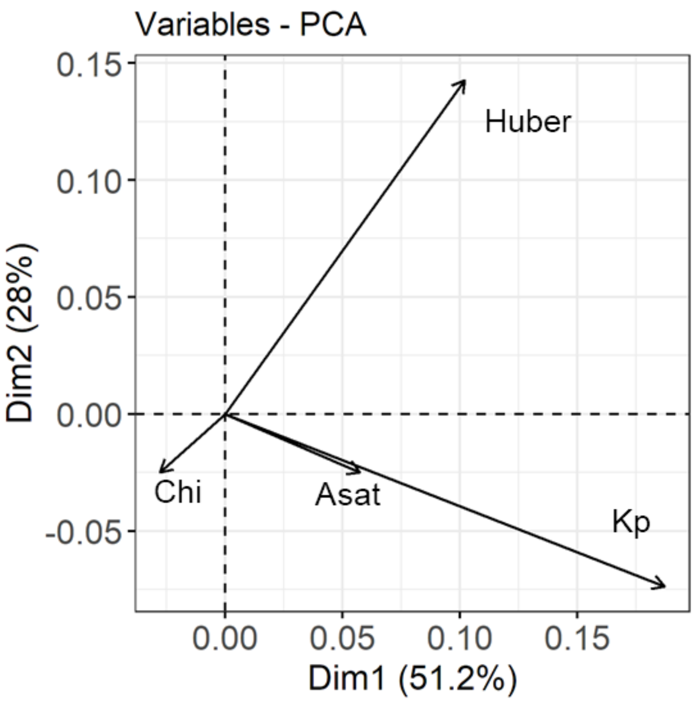

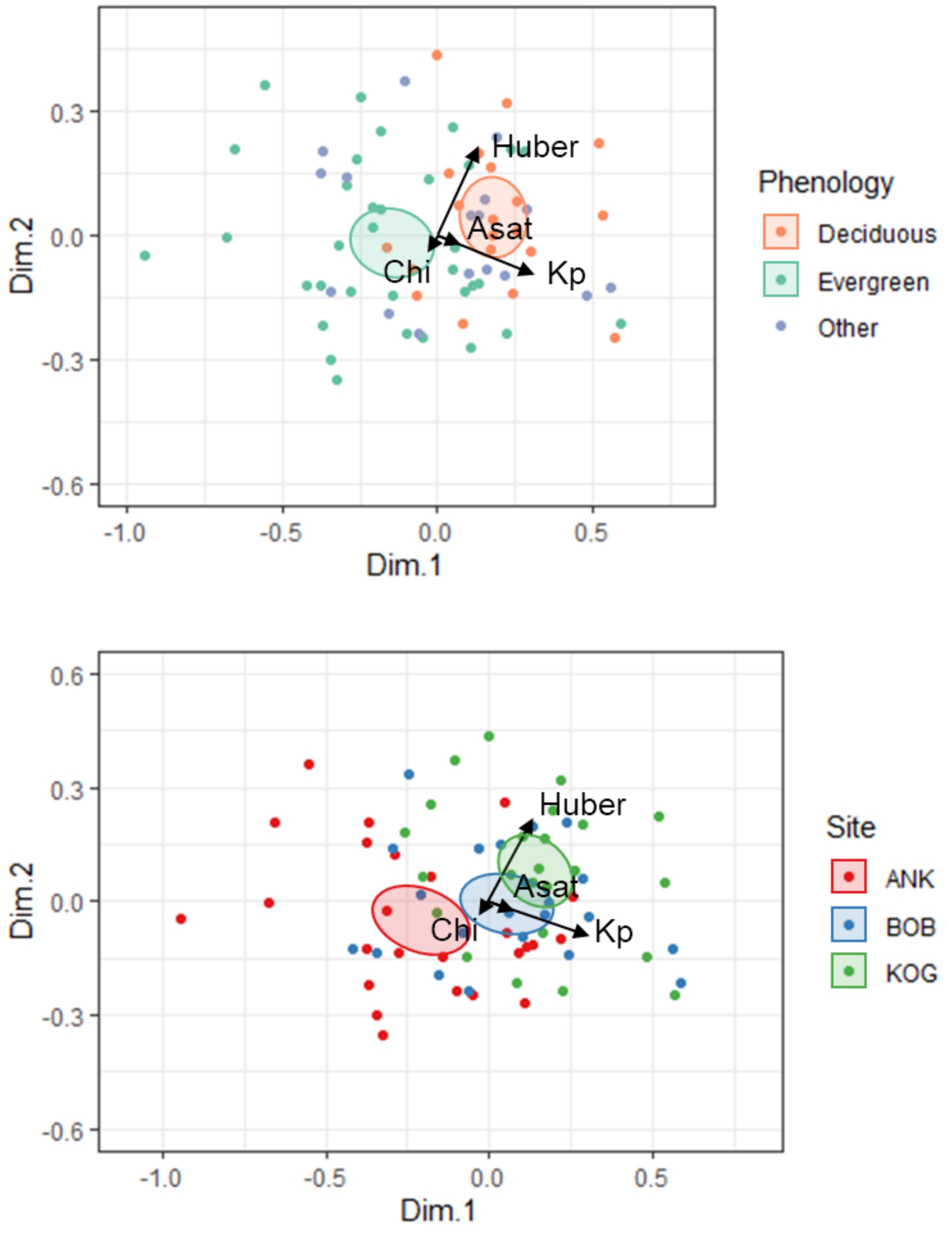


*Figure S 7 Principal components analysis for AS/AL , ci/ca, Asat and Kp. Pleaser also see Figure 2*

#### Appendix 5 Trait extrapolation and Variance partitioning

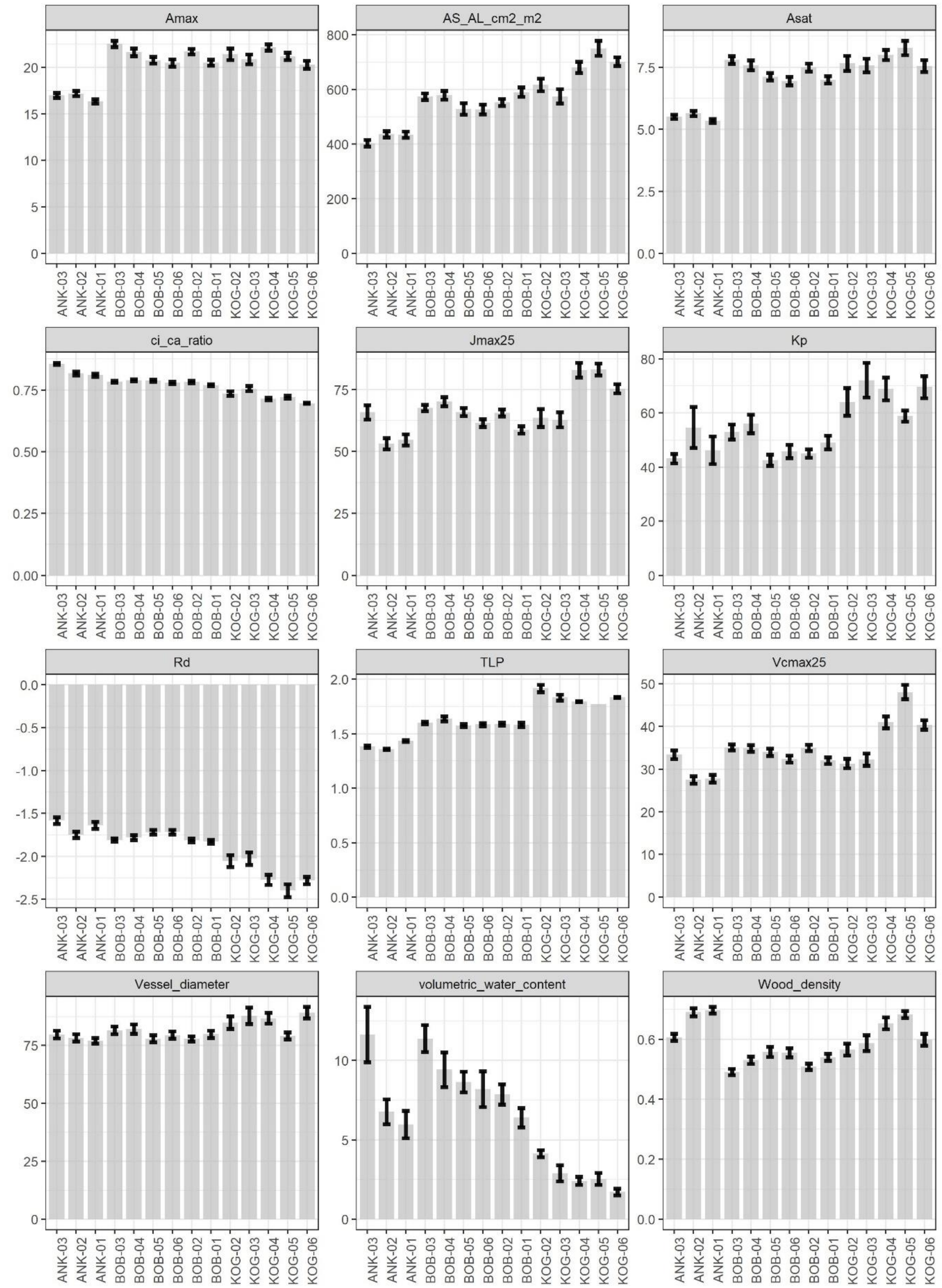


*Figure S 7 Same as Figure 1 but here we extrapolate traits to more sites by assuming the same species share the same trait value, which removes intraspecific variation. The above figure tells the same story as Figure one and hence we conclude that the trait variation pattern (as*

*summarized in Table 1) along the aridity gradient was driven by changing-species instead of intra-specific variation.*

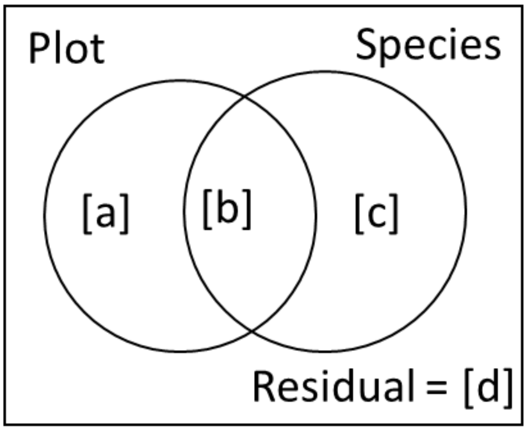


[a] Variance incurred by changing plot for the same species

[b] Variance incurred by changing plot and changing species

[c] Variance incurred by changing Species but not plot (inter-specific variance in a plot)

[d] Variance incurred by anything else but not changing plot and not changing species. It includes intra-specific variation (e.g., changing measurement leaves) and measurement error because for most of the traits, [d] exist only when multiple measurements were collected for one species. If the changing species induced variance [c] is larger than [d], it is safe to conclude that the variation of trait (overall in Ghana) is dominated by changing species, instead of intra-specific variation

**Twig density (g cm-3)**

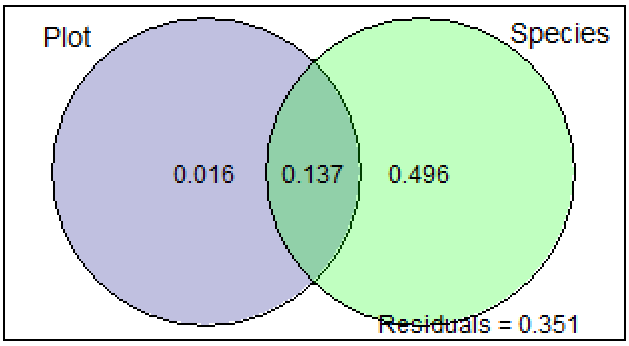


**Vcmax25(umol CO2 m-2 s-1)**

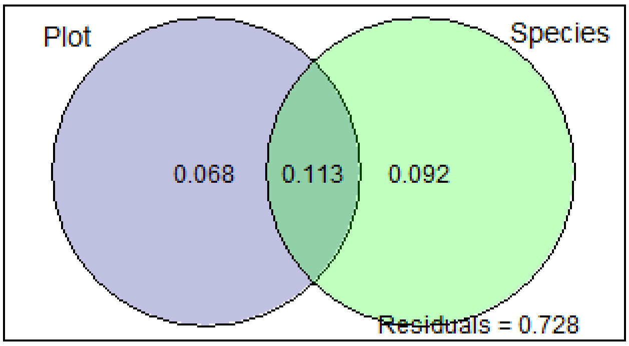


**Vessel diameter (micron)**

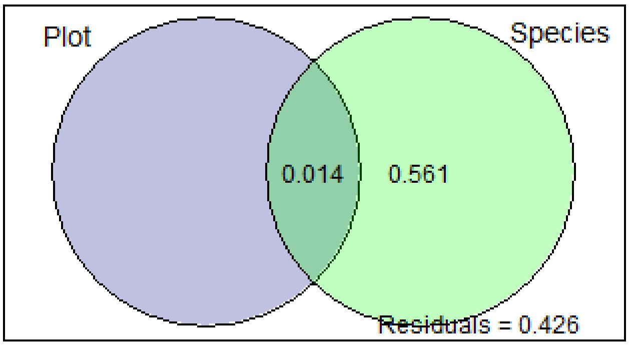


Values <0 not shown

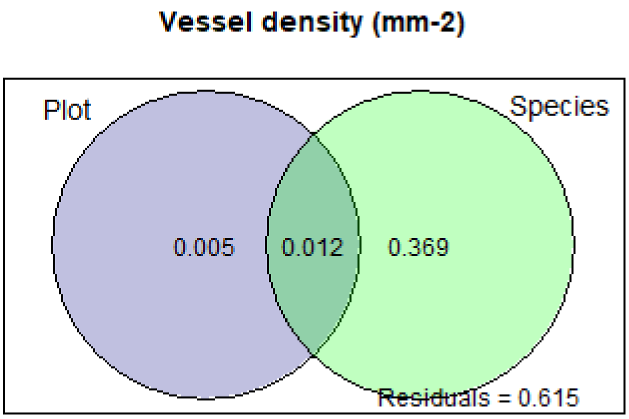
Vessel density (mm-2)
Plot
Species
0.005
0.012
0.369
Residuals = 0.615

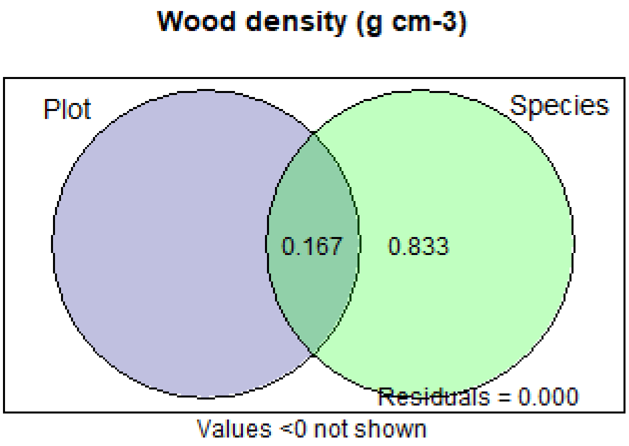
Wood density (g cm-3)
Plot
Species
0.167
0.833
Residuals = 0.000
Values <0 not shown

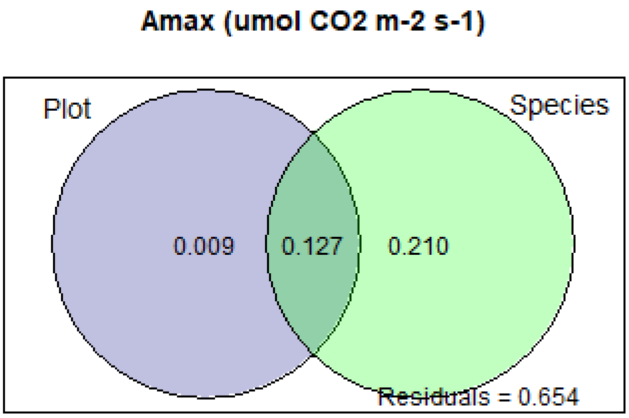
Amax (umol CO2 m-2 s-1)
Plot
Species
0.009
0.127
0.210
Residuals = 0.654

**Asat (umol CO2 m-2 s-1)**

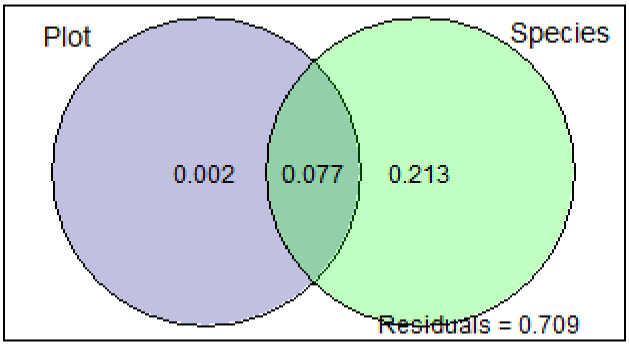


**ci/ca (unitless)**

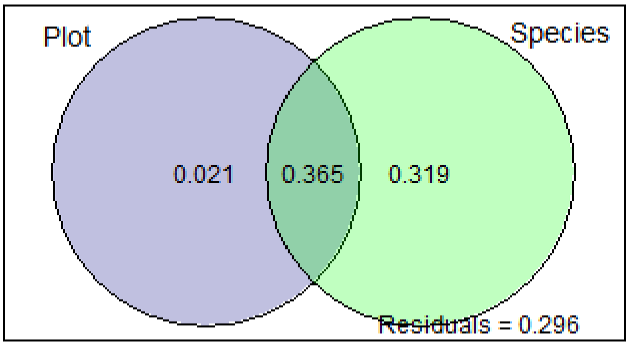


**Rd (umol CO2 m-2 s-1)**

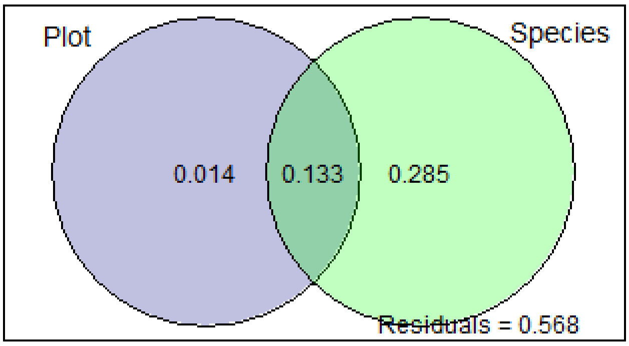

**AS/AL(cm-2 m-2)**

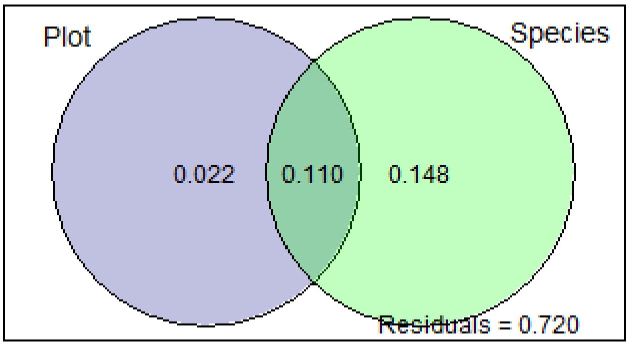


**Jmax25 (umol CO2 m-2 s-1)**

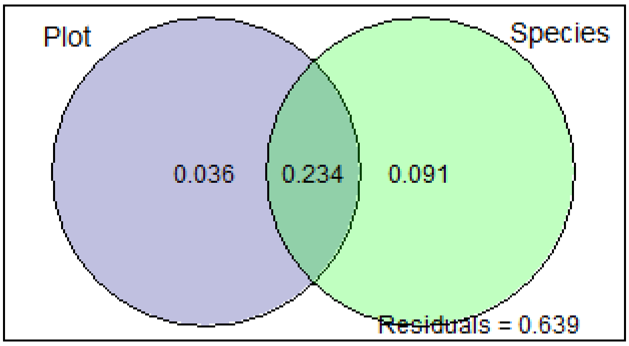


**Kp (kg m-1 MPa-1 s-1)**

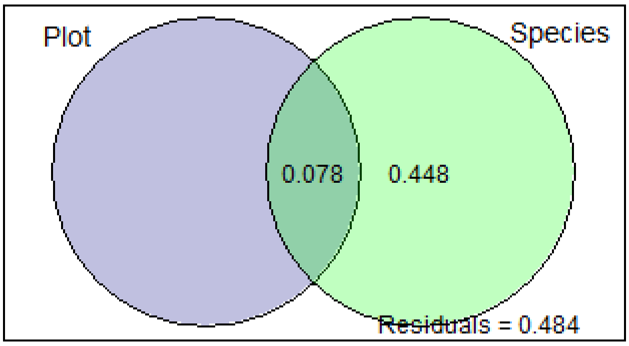


Values <0 not shown

**LMA (g m-2)**

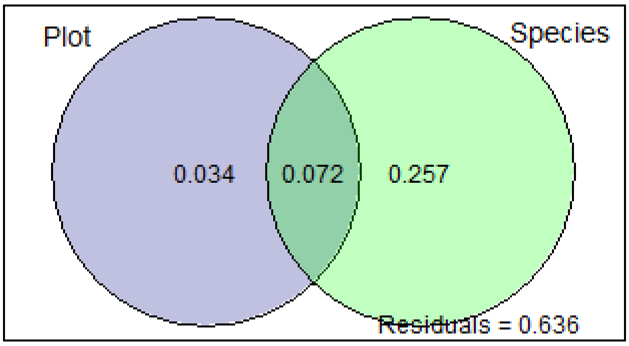


**Narea (g m-2)**

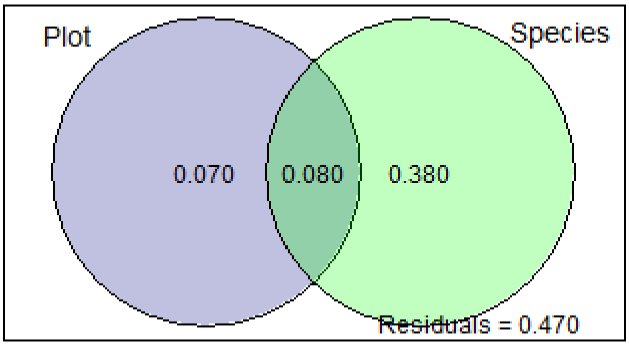


**Nmass (g/kg)**

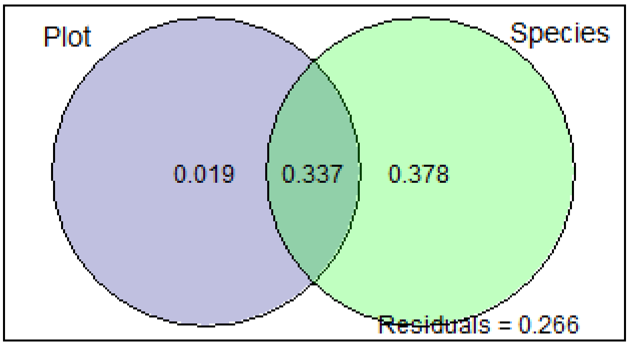

**Parea (g m-2)**

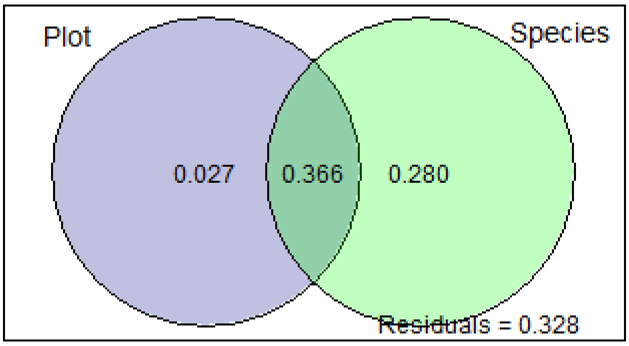


**Pmass (g/kg)**

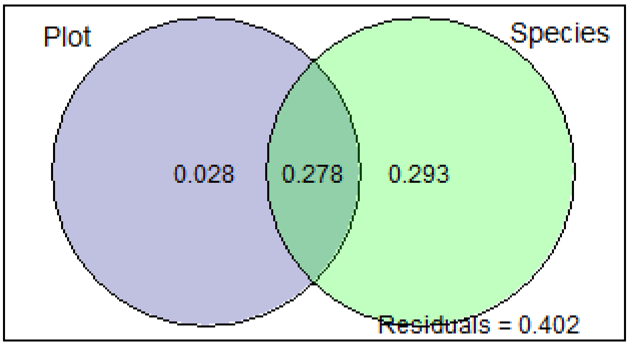


**Rstem_stem (umol m-2 s-1)**

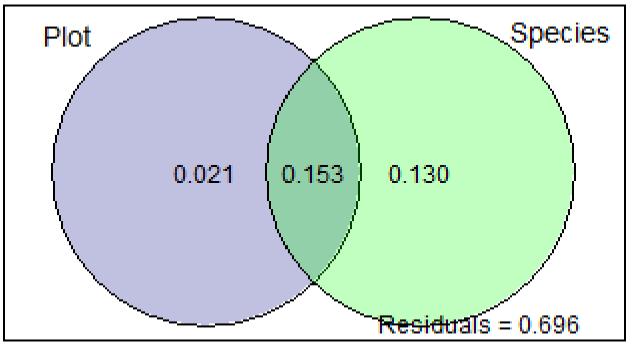

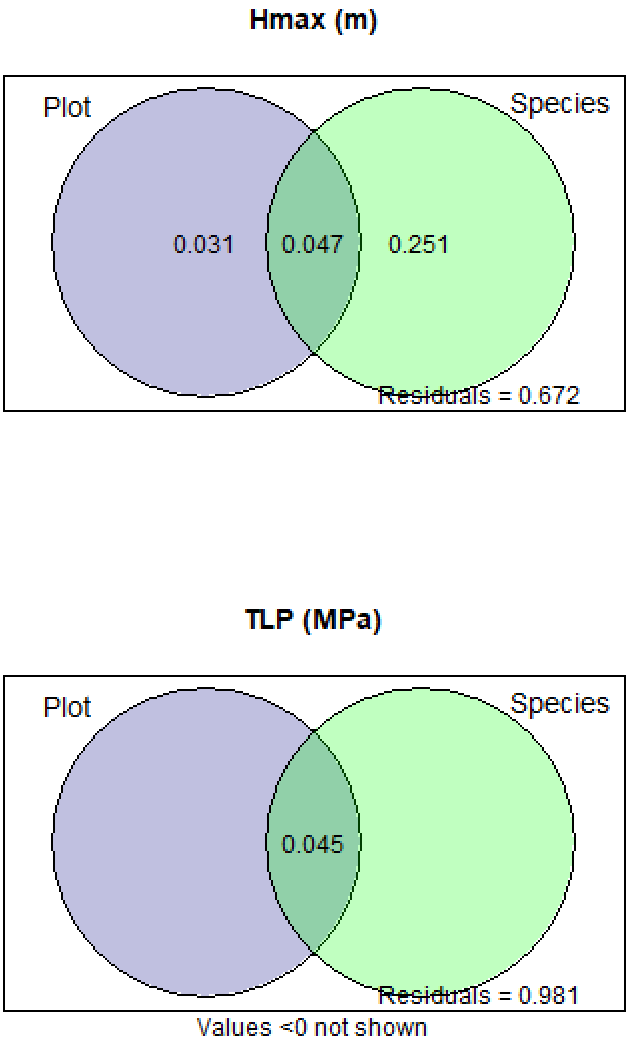


*Figure S 8 Variance partitioning into plot and species. Please see table 1 for definition of traits. Meanings of each number in the circles are explained in the top panel. Note that Vcmax, LMA Narea and Parea from the same plots were published in (Agne 2018), and Asat Amax LMA, Nmass and Pmass from the same plots were published in (Oliveras et al., 2020). Values are not mathematically identical due to (1) different methods of variance partitioning and (2) one more year sampling than the previous publications*

**Aguirre-Gutiérrez J, Oliveras I, Rifai S, Fauset S, Adu-Bredu S, Affum-Baffoe K, Baker TR, Feldpausch TR, Gvozdevaite A, Hubau W, *et al.* 2019**. Drier tropical forests are susceptible to functional changes in response to a long-term drought. *Ecology Letters* **22**: 855–865.

**Appiah D, Osman B, Boafo J**. **2014**. Land Use and Misuse; Human Appropriation of Land Ecosystems Services in Ghana. *International Journal of Ecosystem* **4**: 24–33.

**Choat B, Jansen S, Brodribb TJ, Cochard H, Delzon S, Bhaskar R, Bucci SJ, Feild TS, Gleason SM, Hacke UG, *et al.* 2012**. Global convergence in the vulnerability of forests to drought. *Nature 2012 491:7426* **491**: 752–755.

**Cornwell WK, Wright I, Turner J, Maire V, Barbour M, Cernusak L, Dawson T, Ellsworth D, Farquhar G, Griffiths H**. **2016**. A global dataset of leaf delta 13C values. *Scientific Data*.

**Dong N, Prentice IC, Evans BJ, Caddy-Retalic S, Lowe AJ, Wright IJ, Colin Prentice I, Evans BJ, Caddy-Retalic S, Lowe AJ, *et al.* 2017**. Leaf nitrogen from first principles: Field evidence for adaptive variation with climate. *Biogeosciences* **14**: 481–495.

**Dong N, Prentice IC, Wright IJ, Evans BJ, Togashi HF, Caddy-Retalic S, McInerney FA, Sparrow B, Leitch E, Lowe AJ**. **2020**. Components of leaf-trait variation along environmental gradients. *New Phytologist* **228**: 82–94.

**Gvozdevaite A**. **2018**. The role of economic, venation and morphological leaf traits in plant and ecosystem function along forest-savanna gradients in the tropics.

**Gvozdevaite A, Oliveras I, Domingues TF, Peprah T, Boakye M, Afriyie L, da Silva Peixoto K, de Farias J, Almeida de Oliveira E, Almeida Farias CC, *et al.* 2018**. Leaf-level photosynthetic capacity dynamics in relation to soil and foliar nutrients along forest–savanna boundaries in Ghana and Brazil. *Tree Physiology* **38**: 1912–1925.

**Maire V, Wright IJ, Prentice IC, Batjes NH, Bhaskar R, van Bodegom PM, Cornwell WK, Ellsworth D, Niinemets Ü, Ordonez A**. **2015**. Global effects of soil and climate on leaf photosynthetic traits and rates. *Global Ecology and Biogeography* **24**: 706–717.

**Mencuccini M, Rosas T, Rowland L, Choat B, Cornelissen H, Jansen S, Kramer K, Lapenis A, Manzoni S, Niinemets Ü, *et al.* 2019**. Leaf economics and plant hydraulics drive leaf : wood area ratios. *New Phytologist* **224**: 1544–1556.

**Moore S, Adu-Bredu S, Duah-Gyamfi A, Addo-Danso SD, Ibrahim F, Mbou AT, de Grandcourt A, Valentini R, Nicolini G, Djagbletey G, *et al.* 2018**. Forest biomass, productivity and carbon cycling along a rainfall gradient in West Africa. *Global Change Biology* **24**: e496–e510.

**Niinemets Ü, Wright IJ, Evans JR**. **2009**. Leaf mesophyll diffusion conductance in 35 Australian sclerophylls covering a broad range of foliage structural and physiological variation. *Journal of Experimental Botany* **60**: 2433–2449.

**Oliveras I, Bentley L, Fyllas NM, Gvozdevaite A, Shenkin AF, Peprah T, Morandi P, Peixoto KS, Boakye M, Adu-Bredu S, *et al.* 2020**. The Influence of Taxonomy and Environment on Leaf Trait Variation Along Tropical Abiotic Gradients. *Frontiers in Forests and Global Change* **3**: 18.

**Peng Y, Bloomfield KJ, Prentice IC**. **2020**. A theory of plant function helps to explain leaf-trait and productivity responses to elevation. *New Phytologist* **226**: 1274–1284.

**Poorter L, McDonald I, Alarcón A, Fichtler E, Licona JC, Peña-Claros M, Sterck F, Villegas Z, Sass-Klaassen U**. **2010**. The importance of wood traits and hydraulic conductance for the performance and life history strategies of 42 rainforest tree species. *New Phytologist*.

**Raab N**. **2020**. Non-structural carbohydrates and leaf ecophysiology in tropical and temperate forests.

**Walker AP, Beckerman AP, Gu L, Kattge J, Cernusak LA, Domingues TF, Scales JC, Wohlfahrt G, Wullschleger SD, Woodward FI**. **2014**. The relationship of leaf photosynthetic traits – Vcmax and Jmax – to leaf nitrogen, leaf phosphorus, and specific leaf area: a meta-analysis and modeling study. *Ecology and Evolution* **4**: 3218–3235.

**Wright IJ, Reich PB, Cornelissen JHC, Falster DS, Groom PK, Hikosaka K, Lee W, Lusk CH, Niinemets Ü, Oleksyn J**. **2005**. Modulation of leaf economic traits and trait relationships by climate. *Global Ecology and Biogeography* **14**: 411–421.

**Wright IJ, Reich PB, Westoby M, Ackerly DD, Baruch Z, Bongers F, Cavender-Bares J, Chapin T, Cornellssen JHC, Diemer M, *et al.* 2004**. The worldwide leaf economics spectrum. *Nature 2004 428:6985* **428**: 821–827.

**Zanne AE, Lopez-Gonzalez G, Coomes DA, Ilic J, Jansen S, Lewis SL, Miller RB, Swenson NG, Wiemann MC, Chave J**. **2009**. Data from: Towards a worldwide wood economics spectrum.
